# Perception and memory have distinct spatial tuning properties in human visual cortex

**DOI:** 10.1101/811331

**Authors:** Serra E. Favila, Brice A. Kuhl, Jonathan Winawer

## Abstract

Reactivation of earlier perceptual activity is thought to underlie long-term memory recall. Despite evidence for this view, it is unknown whether mnemonic activity exhibits the same tuning properties as feedforward perceptual activity. Here, we leveraged population receptive field models to parameterize fMRI activity in human visual cortex during spatial memory retrieval. Though retinotopic organization was present during both perception and memory, large systematic differences in tuning were also evident. Notably, whereas there was a three-fold decline in spatial precision from early to late visual areas during perception, this property was entirely abolished during memory retrieval. This difference could not be explained by reduced signal-to-noise or poor performance on memory trials. Instead, by simulating top-down activity in a network model of cortex, we demonstrate that this property is well-explained by the hierarchical structure of the visual system. Our results provide insight into the computational constraints governing memory reactivation in sensory cortex.

## Introduction

Episodic memory retrieval allows humans to bring to mind the details of a previous experience. This process is hypothesized to involve reactivating sensory activity that was evoked during the initial event (James, 1890; Hebb, 1968; Damasio, 1989; McClelland et al., 1995). For example, remembering a friend’s face is thought to involve reactivating neural activity that was present when seeing that face. There is considerable evidence from human neuroimaging demonstrating that the same visual cortical areas active during perception are also active during imagery and long-term memory retrieval (Kosslyn et al., 1995; O’Craven & Kanwisher, 2000; Wheeler et al., 2000; Slotnick et al., 2005; Polyn et al., 2005; Kuhl et al., 2011; Bosch et al., 2014; Waldhauser et al., 2016; Lee et al., 2018; Bone et al., 2018). These studies have found that mnemonic activity in early visual areas like V1 reflects the low-level visual features of remembered stimuli, such as spatial location and orientation (Kosslyn et al., 1995; Thirion et al., 2006; Bosch et al., 2014; Naselaris et al., 2015; Sutterer et al., 2019). Likewise, category-selective activity in high-level visual areas like FFA and PPA is observed when subjects remember or imagine faces and houses (O’Craven & Kanwisher, 2000; Polyn et al., 2005). The strength and pattern of visual cortex activity has been associated with retrieval success in memory tasks (Kuhl et al., 2011, 2013; Gordon et al., 2014), suggesting that cortical reactivation is relevant for behavior.

These studies, and many others, have established similarities between the neural substrates of visual perception and visual memory. However, relatively less attention has been paid to identifying and explaining *differences* between activity patterns evoked during perception and memory. In the present work, we asked the following question: which properties of stimulus-driven activity are reproduced in visual cortex during memory retrieval and which are not? The extreme possibility–that all neurons in the visual system produce identical responses when perceiving vs remembering a given stimulus–can likely be rejected. Early studies demonstrated that sensory responses were reduced during memory retrieval relative to perception (Wheeler et al., 2000), and perception and memory give rise to distinct subjective experiences. A more plausible proposal is that visual memory functions as a “weak” version of feedforward perception (Pearson et al., 2015; Pearson, 2019), with memory activity organized in the same fundamental way as perceptual activity, but with reduced signal-to-noise. This hypothesis is consistent with informal comparisons between perception and memory BOLD amplitudes and data suggesting that visual imagery produces similar behavioral effects to weak physical stimuli in many tasks (Ishai & Sagi, 1995; Pearson et al., 2008; Tartaglia et al., 2009; Winawer et al., 2010). A third possibility is that memory reactivation differs from stimulus-driven activation in predictable and systematic ways beyond signal-to-noise. Such differences could arise due to a change in the neural populations recruited, a change in those populations’ response properties, or a systematic loss of information during sensory encoding or post-sensory processing.

One way to adjudicate between these possibilities is to make use of models from visual neuroscience that quantitatively parameterize the relationship between stimulus properties and the BOLD response. In the spatial domain, population receptive field models (pRF) define a 2D receptive field that transforms stimulus position on the retina to a voxel’s BOLD response (Dumoulin & Wandell, 2008; Wandell & Winawer, 2015). These models are based on well-understood physiological properties of the primate visual system and account for a large amount of variance in the BOLD signal observed in human visual cortex during perception (Kay et al., 2013b). Using these models to quantify memory-evoked activity in the visual system offers the opportunity to precisely model the properties of memory reactivation in visual cortex and their relationship to visual activation. In particular, the fact that pRF models describe neural activity in terms of stimulus properties may aid in interpreting differences between perception and memory activity patterns by projecting these differences onto a small number of interpretable physical dimensions.

Here, we used pRF models to characterize the spatial tuning properties of mnemonic activity in human visual cortex. We first trained human subjects to associate spatially localized stimuli with colored fixation cues. We then measured stimulus-evoked and memory-evoked activity in visual cortex using fMRI. Separately, we fit pRF models to independent fMRI data, which allowed us to estimate receptive field location and size within multiple visual field maps for each subject. Using pRF-based analyses, we quantified the location, amplitude, and precision of neural activity within these visual field maps during perception and memory retrieval. Finally, we explored the cortical computations that could account for our observations by simulating neural responses using a stimulus-referred pRF model and a hierarchical model of neocortex.

## Results

### Behavior

Prior to being scanned, subjects participated in a behavioral training session. During this session, subjects learned to associate four colored fixation dot cues with four stimuli. The four stimuli were unique radial frequency patterns presented at 45, 135, 225, or 315 degrees of polar angle and 2 degrees of eccentricity (Fig. 1a,b). Subjects alternated between study and test blocks (Fig. 1c). During study blocks, subjects were presented with the associations. During test blocks, subjects were presented with the cues and had to detect the associated stimulus pattern and polar angle location among similar lures (Fig. 1a,c; see Methods). All subjects completed a minimum of 4 test blocks (mean = 4.33, range = 4–5), and continued the task until they reached 95% accuracy. Subjects’ overall performance improved over the course of training session (Fig. 1d). In particular, subjects showed improvements in the ability to reject similar lures from the first to the last test block (Fig. 1e).

**Figure 1.**
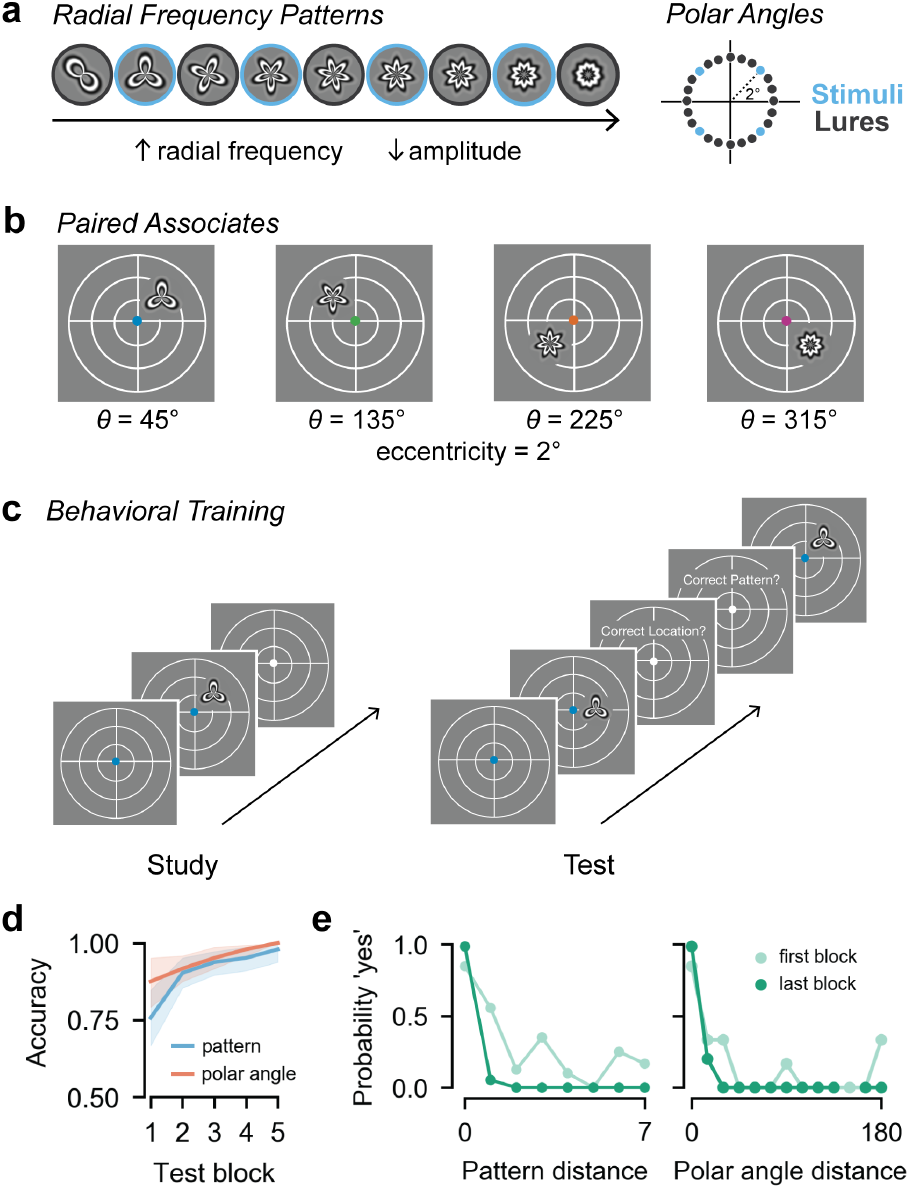
Stimuli and behavioral training. **(a)** The four radial frequency patterns and polar angle locations used in the fMRI experiment are outlined in blue. The intervening patterns and locations were used as lures during the behavioral training session. **(b)** Immediately prior to the scan, subjects learned that each of four colored fixation dot cues was associated with a unique radial frequency pattern that appeared at a unique location in the visual field. **(c)** During training, subjects alternated between study and test blocks. During study blocks, subjects were presented with the associations while maintaining central fixation. During test blocks, subjects were presented with the cues followed by test probes while maintaining central fixation. Subjects gave yes/no responses to whether the test probe was presented at the target polar angle and whether it was the target pattern. **(d)** Accuracy of pattern and polar angle responses improved over the course of the training session. Lines indicate average accuracy across subjects. Shaded region indicates 95% confidence interval. **(e)** Memory performance became more precise from the first to the last test block. During the first block, false alarms were high for stimuli similar to the target. These instances decreased by the last test block. Dots indicates probability of a ‘yes’ response for all trials and subjects in either the first or last block. The x axis is organized such that zero corresponds to the target and increasing values correspond to lures more dissimilar to the target.

After subjects completed the behavioral training session, we collected fMRI data while subjects viewed and recalled the stimuli (Fig. 2a). During fMRI perception runs, subjects fixated on the central fixation dot cues and viewed the four stimuli in their learned spatial locations. Subjects performed a one-back task to encourage covert attention to the stimuli. Subjects were highly accurate at detecting repeated stimuli (mean = 86.9%, range = 79.4%–93.2%). During fMRI memory runs, subjects fixated on the central fixation dot cues and recalled the associated stimuli in their spatial locations. On each trial, subjects made a judgment about the subjective vividness of their memory. Subjects reported that they experienced vivid memory on an average of 89.8% of trials (range: 72.4%–99.5%), weak memory on 8.9% of trials (0.5%–25.0%), and no memory on 1.2% of trials (0.5%–2.6%).

**Figure 2.**
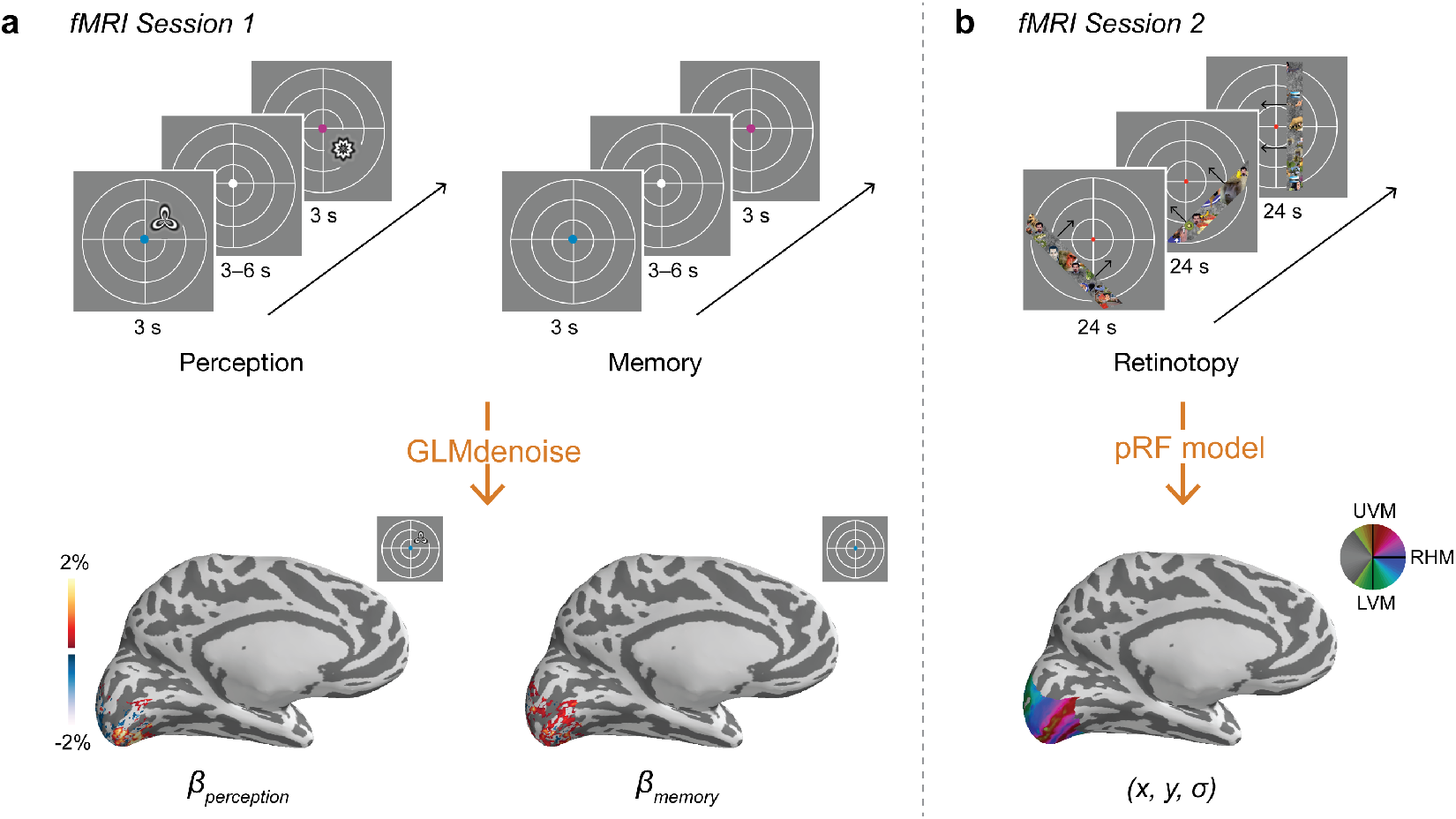
fMRI task design and measurements. **(a)** Following training, subjects participated in two tasks while being scanned. During perception runs, subjects viewed the colored fixation dot cues and associated stimuli while maintaining central fixation. Subjects performed a one-back task on the stimuli to encourage covert attention to each stimulus. During memory runs, subjects viewed only the cues and recalled the associated stimuli while maintaining central fixation. Subjects made a judgment about the vividness of their memory (vivid, weak, no memory) on each trial. We used the perception and memory fMRI time series to perform a GLM analysis that estimated the response evoked by perceiving and remembering each stimulus for each vertex on the cortical surface. Responses in visual cortex for an example subject and stimulus are shown at bottom. **(b)** In a separate fMRI session on a different day, subjects participated in a retinotopic mapping session. During retinotopy runs, subjects viewed bar apertures embedded with faces, scenes, and objects drifting across the visual field while they maintained central fixation. Subjects performed a color change detection task on the fixation dot. We used the retinotopy fMRI time series to solve a pRF model that estimated the receptive field parameters for each vertex on the cortical surface. A polar angle map is plotted for an example subject at bottom.

### Memory reactivation is spatially organized

We used a GLM to estimate the BOLD response evoked by seeing and remembering each of the four spatially localized stimuli (Fig. 2a; see Methods). Separately, each subject participated in a retinotopic mapping session. We fit pRF models to these data to estimate pRF locations (*x,y*) and sizes (σ) in multiple visual areas (Fig 2b). To more easily compare perception- and memory-evoked activity across visual areas, we transformed these responses from cortical surface coordinates into visual field coordinates using the pRF parameters. For each subject, ROI, and stimulus, we plotted the amplitude of the evoked response at the visual field position (*x,*y) estimated by the pRF model (Fig. 3a). We then interpolated these values over 2D space, z-scored the values, rotated all stimulus responses to the same polar angle, and averaged across stimuli and subjects (see Methods). These plots are useful for comparison across regions because they show the organization of the BOLD response in a common space that is undistorted by the size and magnification differences present in cortex.

**Figure 3.**
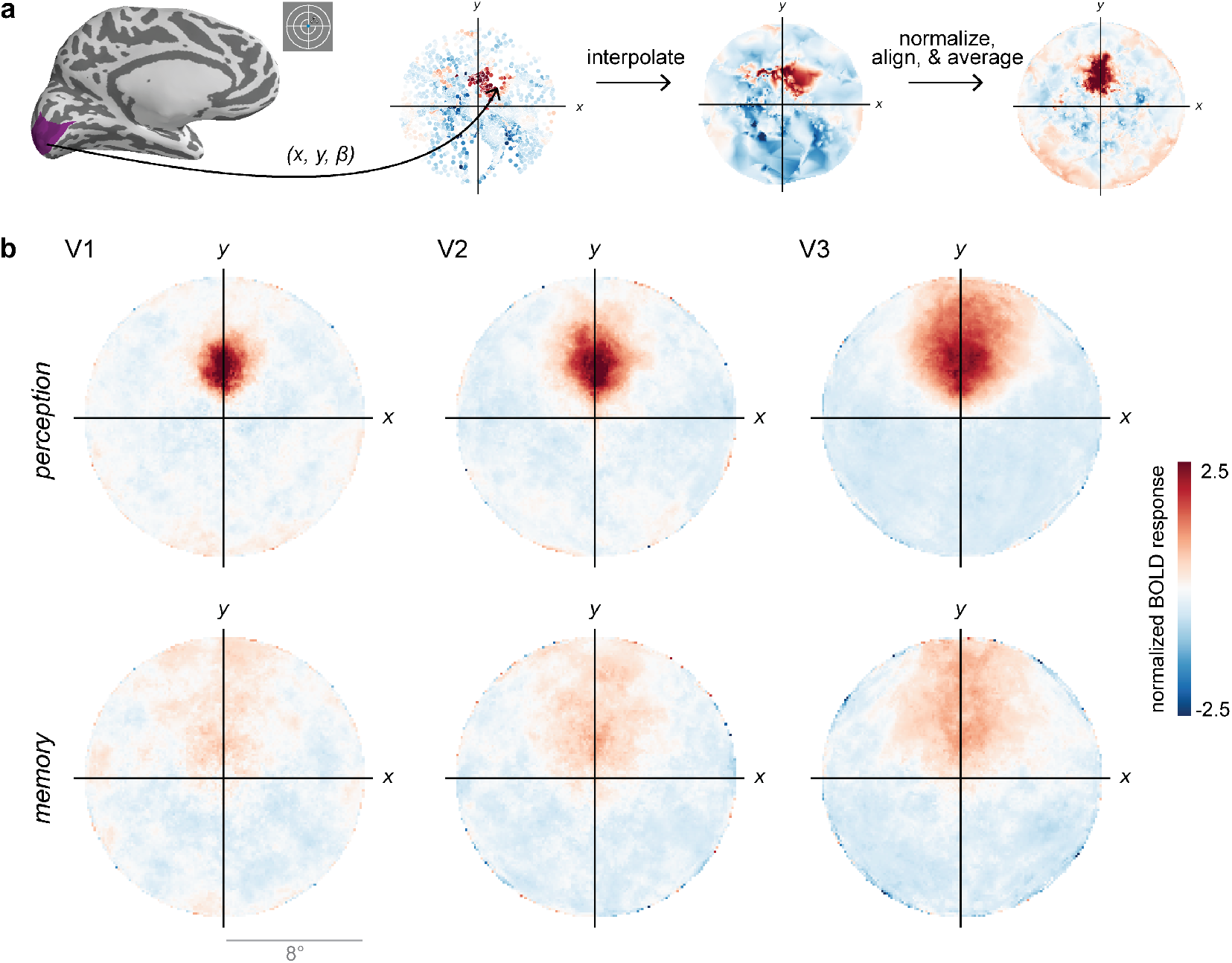
Perception and memory activity in visual field coordinates. **(a)** For a given subject, ROI, and stimulus, we plotted the perception- or memory-evoked response (*β*) in the visual field position estimated by the pRF model *(x,y)*. We then interpolated over 2D space and z-scored the responses. We rotated these representations by the polar angle location of the stimulus so that they aligned on the upper vertical meridian, and then averaged over stimuli. This procedure produces an average activation map in visual field coordinates for each ROI and subject. This map is plotted for V1 in an example subject, at right. **(b)** Plots of perception-evoked and memory-evoked activity, averaged across all subjects, from V1, V2, and V3. These plots reproduce known features of spatial processing during perception, such as increasing receptive field size from V1–V3. They also qualitatively demonstrate that perceptual activity is not perfectly reproduced during memory retrieval but that some retinotopic organization is preserved.

We generated these visual field plots for V1, V2, and V3 as an initial way to visualize the evoked responses during perception and memory. Readily apparent is the fact that stimulus-evoked responses during perception were robust and spatially-specific (Fig. 3b, top). The spatial spread of perceptual responses increased from V1 to V3, consistent with estimates of increasing receptive field size in these regions (Wandell & Winawer, 2015; Kay et al., 2013b). While the memory responses were weaker and more diffuse, they were also spatially organized, with peak activity in the same location as the perception responses (Fig. 3b, bottom).

We quantified these initial observations. Because our stimulus locations were isoeccentric, we reduced our responses to variance along one spatial dimension: polar angle. To do this, we restricted our ROIs to surface vertices with pRF locations near the stimulus eccentricity, rotated stimuli to a common polar angle, normalized the responses, and averaged across stimuli and subjects (see Methods). We then plotted the group average BOLD response in bins of polar angle distance from the stimulus (Fig. 4a). We generated these polar angle response functions for V1–V3 and for three mid-level visual areas: hV4, LO, and V3ab (Fig. 4b). To capture the pattern of positive and negative BOLD responses we observed, we fit the average data in each ROI with a difference of two von Mises distributions, where both the positive and the negative von Mises were centered at the same location. Visualizing the data and the von Mises fits (Fig. 4b), it’s clear that both perception and memory fits show a peak at 0°, or the true location of the stimulus, in every region.

**Figure 4.**
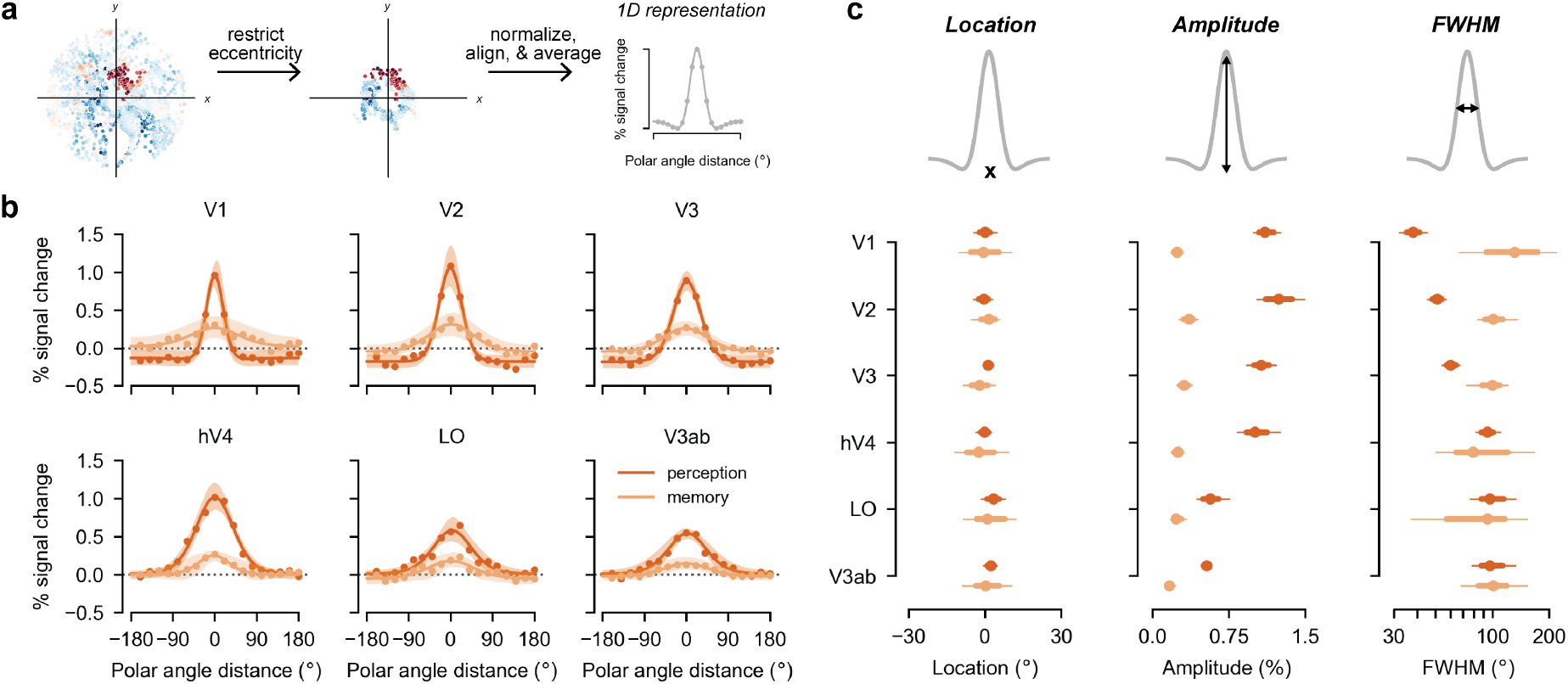
Perception and memory have shared and distinct activation features. **(a)** We created 1D polar angle response functions by restricting data to eccentricities near the stimulus, aligning stimuli to a common polar angle, and averaging responses into polar angle distance bins. A difference of two von Mises distributions was fit to the group average response. Responses in cortical areas that have pRFs near the stimulus position are plotted at x = 0. **(b)** Polar angle response functions, averaged across all subjects and stimuli, are plotted separately for perception and memory. Dots represent average data across all stimuli and subjects. Lines represent the fit of the difference of two von Mises distributions to the average data, and shading represents the 95% confidence interval around this fit. While the peak location of the response is shared across perception and memory, there are clear differences in the amplitude and width of the responses. **(c)** Bootstrapped 68% (thick lines) and 95% confidence intervals (thin lines) for the location, amplitude and FWHM of the difference of von Mises fits are plotted to quantify the responses. In all ROIs, the peak location of the response is equivalent during perception and memory (at 0°, the stimulus location), while the amplitude of the response is reliably lower during memory than during perception. The FWHM of the response increases across ROIs during perception but not during memory, resulting in highly divergent FWHM for perception and memory in early visual areas.

To formally test this, we calculated bootstrapped confidence intervals for the location parameter of the von Mises distributions by resampling subjects with replacement (see Methods). We then compared the accuracy and reliability of location parameters between perception and memory (Fig. 4c, left). As expected, location parameters derived from perception data were highly accurate. 95% confidence intervals for perception location parameters overlapped 0° of polar angle, or the true stimulus location, in all ROIs. These confidence intervals spanned only 7.0° on average (range: 3.9°–9.5°), demonstrating that there was low variability in location accuracy across subjects in every ROI. Critically, memory parameters were also highly accurate, with confidence intervals overlapping 0° in every ROI (Fig. 4c, left). Thus, in every visual area measured, the spatial locations of the remembered stimuli could be accurately estimated from mnemonic activity. Memory confidence intervals spanned 17.6° on average (range = 11.0°–21.3°), indicating that location estimates were somewhat less reliable during memory during perception. However, even the widest memory confidence interval spanned only 21.3°. This is far less than the 90° separating each stimulus location, suggesting that there was no confusability between stimuli present in distributed memory activity. Because both perception and memory location parameters were highly accurate, and because differences in reliability were relatively small, there was no overall difference between perception and memory in the estimated location of peak activity (main effect of perception/memory: *β* = 0.14, 95% CI = [−6.72, 6.14], *p* = 0.94; Fig. 4c, left). These results provide strong evidence that memory-triggered activity in human visual cortex is spatially organized within known visual field maps, as it is during visual perception. These findings support prior reports of retinotopic activity during memory and imagery (Kosslyn et al., 1995; Slotnick et al., 2005; Thirion et al., 2006), but provide more quantitative estimates of this effect.

### Amplitude and precision differ between perceptual and mnemonic activity

Aspects of perception and memory responses other than the peak location differed considerably. First, memory responses were lower in amplitude than perception responses (Fig. 4b). To quantify this observation, we derived a measure of amplitude from the difference of von Mises functions fit to our data (see Methods). We also computed bootstrapped confidence intervals for this amplitude metric, following the prior analysis. We then compared these estimates between perception and memory. First, response amplitudes for perception data were higher than for the memory data (main effect of perception/memory: *β* = 0.95, 95% CI = [0.80, 1.13], p = 0.013; Fig. 4c, middle). The average amplitude during perception was 0.92% signal change, and the average amplitude during memory was 0.26% signal change. Amplitude confidence intervals for perception and memory did not overlap in any ROI, indicating that these differences were highly significant in each region. Critically, the fact the perception amplitudes were larger than memory amplitudes does not imply that memory responses were at baseline. In fact, 95% confidence intervals for memory amplitudes did not overlap with zero in any region (Fig. 4c, middle), demonstrating that responses were significantly above baseline in all areas measured. These results demonstrate that the amplitude of spatially-organized activity in visual cortex is attenuated (but present) during memory retrieval.

Second, memory responses were wider than perception responses (Fig. 4b). We operationalized the *precision* of perception and memory responses by computing the full width at half maximum (FWHM) of the difference of von Mises fit to our data and by generating confidence intervals for this measure. Note that FWHM is not sensitive to the overall scale of the response function: a perception response function rescaled to have the same amplitude as the memory response function will have an unchanged FWHM. On average, FWHM during perception was significantly smaller than during memory (main effect of perception/memory: *β* = −75.2, 95% CI = [−138.5, −33.1], *p* = 0.0002; Fig. 4c, right). However, these differences were not equivalent across ROIs (perception/memory x ROI interaction: *β* = 18.8, 95% CI = [5.78, 35.5], p = 0.021). Specifically, perception FHWM increased moving up the visual hierarchy (main effect of ROI: *β* = 13.3, 95% CI = [10.3, 20.6], p = 0.0056), indicating increased width or *decreased* precision in later visual areas compared to early visual areas (Fig. 4c, right). For example, V1 had the narrowest (most precise) response during perception, with an average FWHM of 38.0°(95% CI: [32.0°, 45.0°]), while V3ab had the widest responses during perception, with a FWHM of 97.0°(95% CI: [78.0°, 132.5°]). This increasing pattern follows previously described increases in population receptive field size in these regions (Wandell & Winawer, 2015; Kay et al., 2013b). Note that a separate question, not addressed here, is the precision with which the stimulus can be decoded from a representation, which is not necessarily related to receptive field size.

Strikingly, this pattern of increasing FWHM from early to late visual areas was abolished during memory (main effect of ROI: *β* = −5.49, 95% CI = [−18.7, 8.41], p = 0.20; Fig. 4c, right). For areas V1–hV4, the regions we can sort hierarchically with the most confidence, the pattern across ROIs trended toward being reversed, with the widest responses observed in the *earliest* areas (main effect of ROI: *β* = −15.7, 95% CI = [−39.8, 12.4], p = 0.083). These data demonstrate that fundamental aspects of spatial processing commonly observed during perception do not generalize to memory-evoked responses. Interestingly, the interaction between perception/memory and ROI yielded highly divergent perception and memory responses in the earliest visual areas but equivalent responses in the latestvisual areas (Fig. 4c, right). For example, V1 responses during memory had an average FWHM of 131.0° (95% CI: [66.9°, 225.0°]), and were thus 3.45 times wider than V1 responses during perception. In V2 and V3, memory FWHM exceeded perception FWHM by an average of 1.98 times and 1.67 times, respectively. Unlike in V1-V3, confidence intervals for perception and memory were highly overlapping in hV4, LO, and V3ab (Fig. 4c, right). In these later areas, memory responses were only 0.84–1.04 times wider during memory than during perception. These data raise the interesting possibility that later stages of perceptual processing serve as a bottleneck on mnemonic activity precision. Taken together, these results provide evidence for reliable and striking differences in the precision of perception and memory activity across different levels of the visual system. More broadly, these findings indicate that there are fundamentally different constraints on the properties of feedforward perceptual activity and top-down mnemonic activity in human visual cortex.

### Differences between perception and memory responses are not explained by noise

One important consideration in interpreting our results is whether the differences we observed between perception and memory could be caused by differences in noise. For example, is it possible that perception and memory responses were actually equivalent other than noise level, but due to greater trial-to-trial noise, memory responses appeared to have systematically different tuning? In particular, we sought to understand whether differences in memory responses could be explained by three types of noise: 1) reduced fMRI signal-to-noise; 2) retrieval failure on a subset of trials; 3) memory error. If perception and memory have the same fundamental response properties, but memory is subject to more noise, then adding noise to the perception data should yield responses that look like what we observed during memory. Thus, we started with perception data (mean and variance of each voxel’s activity during perception) and tested whether we could generate responses that looked like memory data by adding one of the three types of noise. To simulate reduced fMRI signal-to-noise, we introduced additive noise to each voxel’s perception response (Fig. 5a, left; see Methods). To simulate retrieval failure, we created some trials where the mean response was zero (Fig. 5b, left). To simulate memory error, we added angular noise to the peak location of the perception responses (Fig. 5c, left). For each of these types of simulation, we considered multiple levels of noise. To assess the simulation results, we analyzed all simulated datasets with the same procedures used for the real data and then counted the proportion of times the von Mises parameters derived from a simulation fell within the 95% confidence interval of the actual memory data (Fig. 5a-c, right).

**Figure 5.**
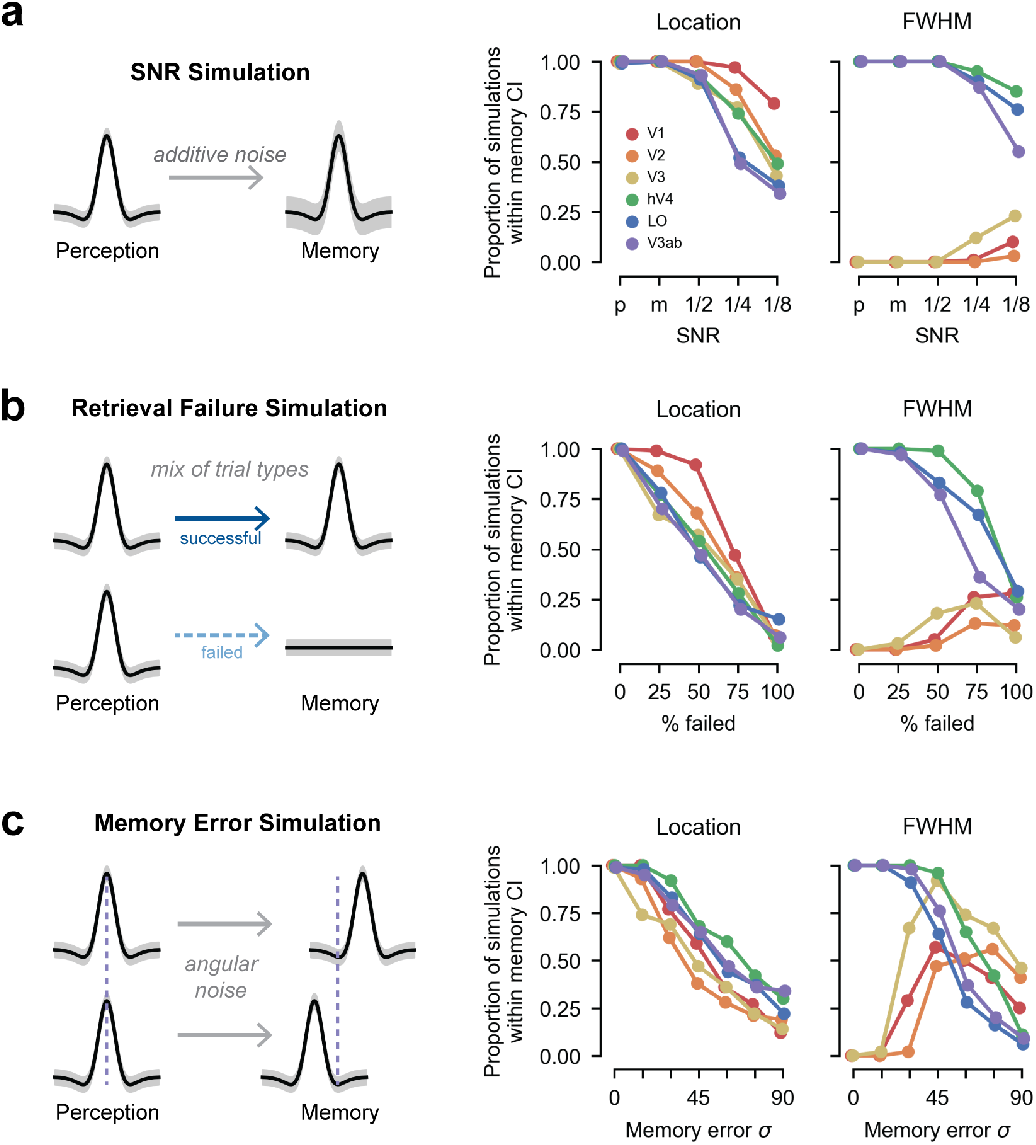
Differences between perception and memory are not explained by noise. **(a)** Left: We simulated the effect of low SNR by introducing additive noise to our perception data and asked whether this was sufficient to produce responses similar to what we observed during memory. Right: Proportion of simulations that produce location and FWHM parameters within the 95% confidence intervals of the memory data are plotted for decreasing signal-to-noise ratios (SNR) and for each ROI. SNR values ranged from the empirical SNR of the perception data (p), the empirical SNR of the memory data (m), or 1/2, 1/4, or 1/8 of the empirical SNR of the memory data. Even at extremely high noise levels, very few simulations generate FWHM parameters within the confidence intervals of the memory data in V1–V3. **(b)** Left: We simulated the effect of failing to perform the retrieval task by generating a perception dataset where a subset of trials had a mean BOLD response of zero. Right: Data are plotted as in a, with increasing large numbers of failed trials on the x axis. As in a, even at extremely high rates of failed retrieval, FWHM parameters in V1–V3 rarely fall within the memory confidence interval **(c)** Left: We simulated the effect of memory error by adding angular noise to the peak location of the perception responses. Right: Data are plotted in a, with increasing large amounts of memory error on the x axis. Implausibly large amounts of memory error are needed to generate FWHM parameters that fall within the memory confidence intervals >= 50% of the time in V1–V3. In all panels, increased noise produced a worse match to memory FWHM in hV4, LO and V3ab, as well as unreliable location parameters in all ROIs.

First, using bootstrapped parameter estimates, we confirmed that the estimated signal-to-noise ratio (SNR) for perception parameter estimates was higher than for memory parameter estimates in every ROI. Perception SNR was between 1.3 and 1.6 times higher than memory SNR in each ROI, and between 2.2 and 4.3 times higher in vertices closest to the stimulus location. Given this, we simulated new perception data that precisely matched the empirical memory SNR for every surface vertex. We also simulated data with even lower SNR (higher noise) than what we observed during memory. As expected, simulating perception data with reduced SNR increased variance in the location, amplitude, and FWHM of the von Mises fits (Supplementary Fig. 1a). However, no level of SNR produced response profiles that matched the memory data well. In V1–the region where we observed the largest difference in FWHM between perception and memory–0% of the FWHM parameters in the memory SNR simulation approximated the actual memory data (Fig. 5a, right). In the noisiest simulation we performed (1/8 of the memory SNR), this figure was still only 10% (Fig. 5a, right). Similar results occurred for V2 and V3. These simulations demonstrate that low SNR cannot explain the pattern of memory responses we observed in early visual cortex.

Our SNR simulations also demonstrate that there are fundamental tradeoffs between capturing memory FWHM in early visual cortex and in capturing other aspects of the data. First, at high levels of noise (low levels of SNR), any modest increase in ability to capture V1-V3 FWHM was accompanied by a *decrease* in ability to capture FWHM in later visual areas (Fig. 5a, right). In these regions, FWHM was already equivalent during perception and memory, and artificially adding noise to the perception data harms this equivalence. Second, high noise simulations generated more noise in the location parameters than was actually observed in the memory data (Fig. 5a, right), resulting in unreliable location parameters in all ROIs.

We observed a similar pattern of results in the retrieval failure simulations. Very high rates of retrieval failure were required to generate any FWHM parameters that were sufficiently wide to match the memory data in V1 (Supplementary Fig. 1b). Only when simulating retrieval failure in >=50% of all trials, did this number exceed 0% (Fig. 5b, right). Similar to the SNR simulation, any improvement in ability to capture the V1 FWHM data with increased retrieval failure was offset by a decline in ability to capture FWHM in late visual areas (Fig. 5b, right), where responses became much wider than what was observed empirically during memory. Again, as in the SNR simulation, high rates of retrieval failure were also associated with location parameters that were far noisier than what we observed (Fig. 5b, right). Thus, subjects experiencing retrieval failure on a subset of trials does not explain our memory data.

Finally, we considered the memory error simulation. Compared to the other simulations, this simulation produced a better match to memory FWHM in V1 when assuming high levels of noise (Supplementary Fig. 1c). Still, in the best performing simulation only 57% of the V1 FWHM parameters approximated the memory data (Fig. 5c, right). Critically, the magnitude of memory error in this simulation was implausibly high. The standard deviation of memory errors around the true value was 45°, meaning that simulated memories were within the correct quadrant only 68% of time. Given that subjects were trained to discriminate remembered locations up to 15° (see Methods), errors of this magnitude and frequency are exceedingly unlikely. At a more plausible 15° standard deviation of memory error, 0% of simulations approximated the memory data (Fig. 5c, right). Further, similar to the other simulations, improvements in the ability to capture V1 FWHM with high levels of angular error were accompanied by decreases in the ability to capture FWHM in later areas and by decreases in location parameter reliability beyond what we observed empirically (Fig. 5c, right). Thus, subjects experiencing a small, variable amount of memory error does explain our memory data.

Collectively, these simulations demonstrate that our results are unlikely to be caused by a simple source of measurement noise (reduced SNR) or cognitive noise (failed retrieval, memory error). In each of the three simulations, the amount of noise required to make even modest gains in our ability to account for the V1 memory FWHM was implausibly large. Further, in all three cases, increases in the ability to account for V1 FWHM were accompanied by decreases in the ability to account for FWHM in higher visual areas and to recover location parameters that were as reliable as our actual data.

### pRF models accurately predict perception but not memory responses

Next, we evaluated how well perception and memory responses matched the predictions of a pRF model. To do this, we used a modified version of each subject’s pRF model to generate predicted cortical responses to each of the four experimental stimuli (Fig. 6a; see Methods). The pRF model we used to generate predictions is a novel variant of existing pRF models: we added a Difference of Gaussian pRF shape (Zuiderbaan et al., 2012) with a fixed positive to negative Gaussian size ratio (1:2) and amplitude ratio (2:1) to our solved nonlinear compressive spatial summation (CSS) model (Kay et al., 2013b). The predictions from the model were analyzed with the same procedure as the data, yielding von Mises fits to the predicted data (Fig. 6b). Model predictions from simpler pRF models are shown in Supplementary Figure 2.

**Figure 6.**
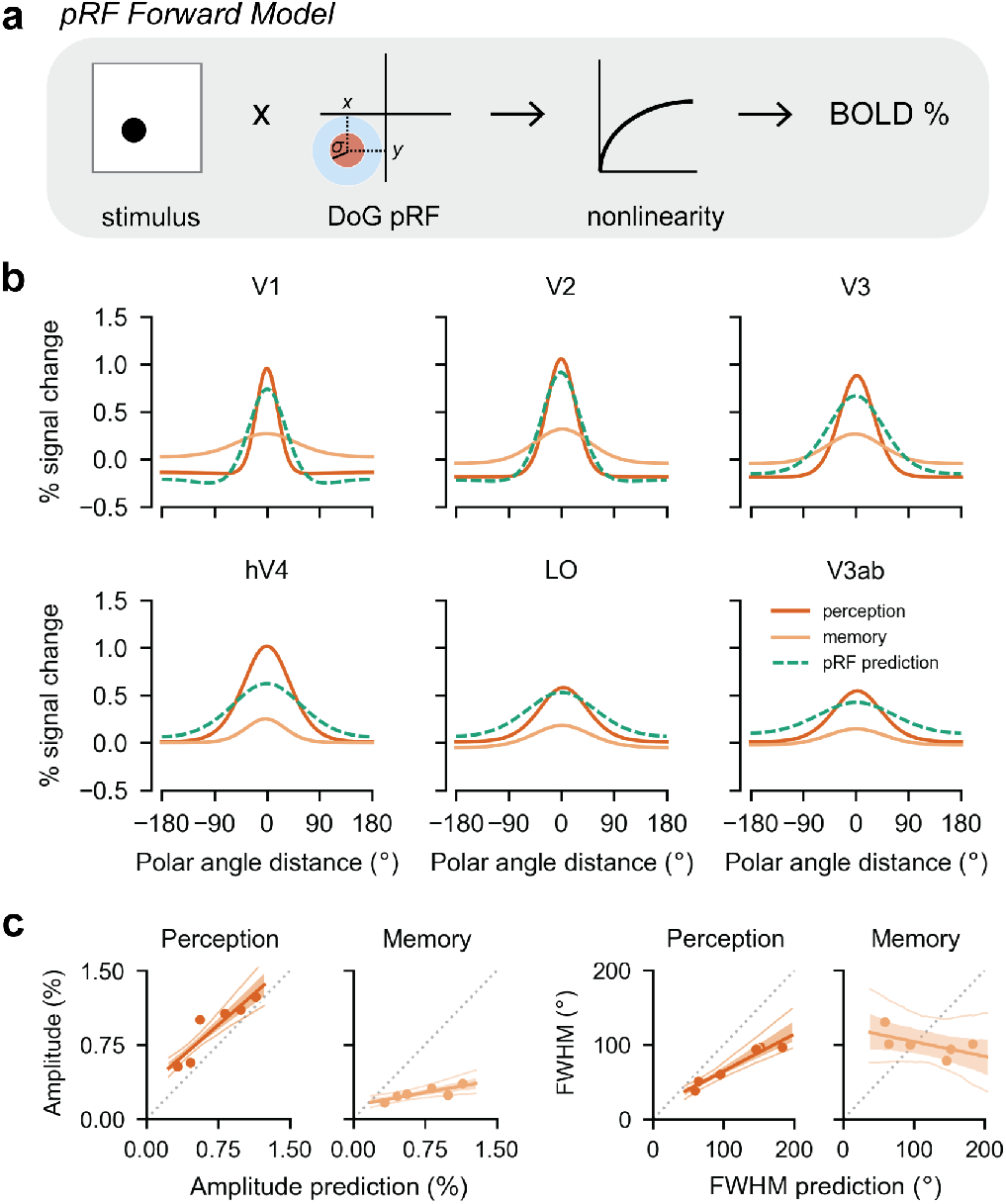
pRF forward model captures perception but not memory responses. **(a)** We used our pRF model to generate the predicted BOLD response to each of our experimental stimuli. The model assumes a Difference of Gaussians pRF shape, with a fixed positive to negative Gaussian size ratio (1:2) and amplitude ratio (2:1). The model also incorporates a compressive nonlinearity, **(b)** Predicted polar angle response functions are plotted for the pRF model (green dashed lines), alongside the functions fit to the perception and memory data (dark and light orange, reproduced from Fig. 4b). The model predictions are closer to the perception data than the memory data in all visual areas. **(c)** Predicted versus observed amplitude (left) and FWHM (right), plotted separately for perception and memory. Each dot represents an ROI. The shaded region is the 68% CI from bootstrapping linear fits across participants, and the thin lines indicate the 95% confidence intervals. For both the amplitude and FWHM, the perception data lie relatively close to the pRF model predictions (dashed grey lines), whereas the memory data do not.

Qualitatively, the pRF model predictions agree with the perception data but not the memory data (Fig. 6b). Several specific features of the perception data are well captured by the model. First, the model predicts the highest amplitude response at cortical sites with pRFs near the stimulus location (peak at 0°). Second, the model predicts increasingly wide response profiles from the early to late visual areas. Third, it predicts higher amplitudes in early compared to late areas. Finally, the model predicts negative responses in the surround locations of V1-V3 but not higher visual areas. This is particularly interesting given that all voxel pRFs were implemented with a negative surround of the same size and amplitude relative to the center Gaussian. This suggests that voxel-level parameters and population-level responses can diverge (Sprague & Serences, 2013, see also). Though not the focus of this analysis, we note that the model predictions are not perfect. The model predicts slightly lower amplitudes and larger FWHM than is observed in the perception data (Fig. 6b). These discrepancies may be due to differences between the stimuli used in the main experiment and those used in the pRF experiment or to differences in the task (attending fixation during the pRF experiment vs attending the stimulus during the main experiment).

Critically, the model accurately captures the properties of memory responses that are shared with perception responses (the peak location), but not the distinct properties (Fig. 6b). These failures are especially clear when comparing the predicted amplitude and FWHM from the pRF model with the observed amplitudes and FWHMs for perception and memory. While there is a positive slope between the predicted amplitude and both the perception (*β* = 0.84, 95% CI: [0.56, 1.15]) and memory amplitudes (*β* = 0.17, 95% CI: [0.056, 0.32]), the slopes differ substantially (Fig. 6c). The perception amplitudes have a slope closer to 1, indicating good agreement with the model predictions, while the memory data have a slope closer to 0, indicating weak agreement. Similarly, the predicted FWHM is strongly and positively related to the perception FWHM (*β* = 0.50, 95% CI: [0.36, 0.76]), but weakly and negatively related to the memory FWHM (*β* = −0.20, 95% CI: [−0.67, 0.26]; Fig. 6c). These analyses strongly support our interpretation of the data in Figure 4b,c to mean that memory and perception have distinct spatial tuning properties. The critical advantage of using pRF models is that they explicitly incorporate known properties of feedforward spatial processing in visual cortex. Because our pRF model fails to account for the memory responses we observed, we can conclude that memory reactivation violates the assumptions of feedforward processes that well characterize perceptual activation. A plausible explanation for this failure is that memory retrieval involves a fundamentally different origin and cascade of information through visual cortex, a possibility we explore in detail in the next section.

### Perception and memory responses can be simulated with a bidirectional hierarchical model

Cortical activity during perception arises from a primarily feedforward process that originates with the retina and that accumulates additional spatial pooling in each cortical area, resulting in increasingly large receptive fields (Gattass et al., 2005; Wandell & Winawer, 2015). In contrast, memory reinstatement is hypothesized to begin with the hippocampus (Marr, 1971; O’Reilly & McClelland, 1994), a region bidirectionally connected to high-level visual areas in ventral temporal cortex via the medial temporal lobe cortex (Van Hoesen & Pandya, 1975; Felleman & Essen, 1991; Suzuki & Amaral, 1994). Reinstated cortical activity is then thought to propagate backwards through visual cortex (Naya et al., 2001; Linde-Domingo et al., 2019; Dijkstra et al., 2019; Hindy et al., 2016). Here, we explored whether a simple hierarchical model with spatial pooling could be adapted to account for both our perception and memory results by manipulating the direction of information flow.

We first constructed a linear feedforward hierarchical model of spatial processing in neocortex. In this model, the activity in each layer was created by convolving the activity from the previous layer with a fixed Gaussian kernel (Fig. 7a; see Methods). Beginning with a boxcar stimulus, we cascaded this convolutional operation to simulate 8 layers of the network (Fig. 7b). In this simple demonstration, the size of the convolutional kernel was fixed, not fit to the data. Nonetheless, the pattern of feedforward responses qualitatively matches our fMRI observations during perception. The location of the peak response is unchanged across layers, but response functions become wider and lower in amplitude in higher layers (Fig. 7b,c)–precisely as we observed in our actual data (Fig. 4b,c).

**Figure 7.**
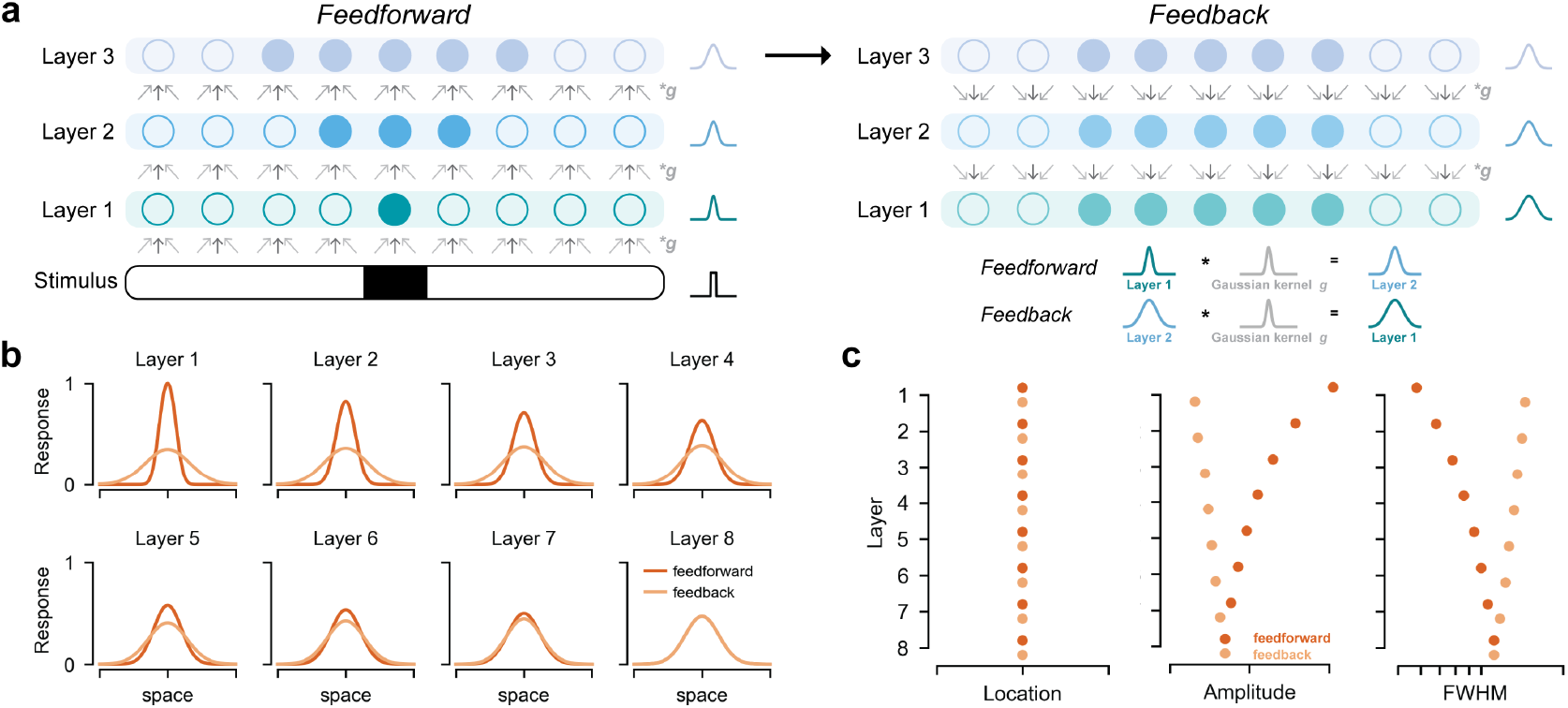
Perception and memory responses can be simulated with a bidirectional hierarchical model. **(a)** Illustration of stimulus-driven activity propagating through a linear hierarchical network model in the feedforward direction (left) and mnemonic activity propagating through the model in the feedback direction (right). In both cases, a given layer’s activity is generated by convolving the previously active layer’s activity with a fixed Gaussian kernel. The feedforward simulation began with a boxcar stimulus. The feedback simulation began with duplication of the feedforward activity from the final layer. **(b)** Results from feedforward and feedback simulations in an 8 layer network, plotted in the conventions of Figure 4b. The feedforward simulation parallels our observations during perception, and the feedback simulation parallels our observations during memory. **(c)** Location, amplitude, and FWHM parameters for each layer, plotted separately for feedforward and feedback simulations. Location is preserved across layers in the feedforward and feedback direction. Note that FWHM become progressively wider in later layers in the feedforward direction and in earlier layers in the feedback direction. This results in large differences in FWHM between feedforward and feedback activity in early layers. These trends closely follow our observations in Figure 4c.

We then explored whether backwards propagation of reinstated activity in our hierarchical model could account for our memory data. To do this, we assumed that feedforward and feedback connections in the model were reciprocal, meaning that the convolutional kernel was the same in feedforward and feedback direction. We assumed perfect reinstatement in the top layer, and thus began the feedback simulation by duplicating the feedforward activity from the final layer. Starting with this final layer activity, we convolved each layer’s activity with the same Gaussian kernel to generate earlier layers’ activity (Fig. 7a). The properties of the simulated activity (Fig. 7b,c) bear a striking resemblance to the those of the observed memory data (Fig. 4b,c). First, simulated feedback activity had a preserved peak location across layers (Fig. 7c, left), similar to the memory data. Second, simulated feedback activity was wider and lower amplitude than feedforward activity overall (Fig. 7c, middle and right)-just as the memory data had wider and lower amplitude responses than the perception data. Third, the increase in FWHM across layers was smaller in the feedback direction than in the feedforward direction, and it reversed direction with respect to the visual hierarchy (Fig. 7c, right). This small effect of reversal is particularly interesting given that this trend was numerically present in our memory data but not statistically reliable at our sample size. Finally, the difference between feedforward and feedback FWHM was maximal in the earliest layers (Fig. 7c, right), just as the difference between our perception and memory data was maximal in V1. This simulation suggests that the distinct spatial profile of mnemonic responses in visual cortex may be a straightforward consequence of reversing the flow of information in a system with hierarchical structure and reciprocal connectivity, and that spatial pooling accumulated during feedforward processing may not be inverted during reinstatement. More broadly, these results demonstrate that models of the visual system may be useful for probing the mechanisms that support and constrain visual memory.

## Discussion

In the current work, we combined empirical and modeling approaches to explore how long-term spatial memories are represented in the human visual system. By using computational models of spatial encoding to compare perceptual and mnemonic BOLD activity, we provide strong evidence that visual memory, like visual perception, produces retinotopically-mapped activation throughout visual cortex. Critically, however, we also identified systematic differences in the population spatial tuning properties of perceptual and mnemonic activity. Compared to perceptual responses, mnemonic responses were lower in amplitude in all visual areas. Further, while we observed a threefold change in spatial precision from early to late visual areas during perception, mnemonic responses violated this pattern. Instead, mnemonic responses displayed consistent spatial precision across visual areas. Notably, simulations showed that neither reduced SNR, nor failure to retrieve on some trials, nor memory error could account for this difference. We speculate, instead, that this difference arises from a reversal of information flow in a hierarchically organized and reciprocally connected visual cortex. To support this, we show that top-down activation in a simple hierarchical model elicits a systematically different pattern of responses than bottom-up activation. These simulations reproduce the properties we observe during both perception and memory. Together, these results reveal novel properties of memory-driven activity in visual cortex that suggest specific computational processes governing visual cortical responses during memory retrieval.

### Advantages of using encoding models to parameterize memory representations

Much work in neuroscience has been dedicated to the question of how internally-generated stimulus representations are coded in the brain. Early neuroimaging work established that sensory cortices are recruited during imagery and memory tasks (Kosslyn et al., 1995; O’Craven & Kanwisher, 2000; Wheeler et al., 2000), moving the field away from purely symbolic accounts of memory (e.g. Pylyshyn, 2002). More recently, memory researchers have favored decoding and pattern similarity approaches over univariate activation analyses to examine the content of retrieved memories (Polyn et al., 2005; Kuhl et al., 2011; Favila et al., 2018). While these approaches are powerful, they do not explicitly specify the form mnemonic activity should take, and many activation schemes can lead to successful decoding or changes in pattern correlations. In the present work, we leveraged encoding models from visual neuroscience, specifically stimulus-referred pRF models, to examine and account for memory-triggered activity in visual cortex. In contrast to decoding or pattern similarity approaches, encoding models predict the activity evoked in single voxels in response to sensory or cognitive manipulations using a set of explicit mathematical operations (Naselaris et al., 2011). Spatial encoding models have proved particularly powerful because space is coded in the human brain at a scale that is well-matched to the millimeter sampling resolution of fMRI (Engel et al., 1994; Sereno et al., 1995; Dougherty et al., 2003). Despite the power of such encoding models, relatively little work has applied these models to questions about long-term memory (c.f. Thirion et al., 2006; Naselaris et al., 2015; Breedlove et al., 2018). Here, using this approach, we revealed novel properties of memory responses in visual cortex that decoding approaches have missed. Most notably, we found that memory activity was characterized by a different pattern of spatial precision across regions than perceptual activity. Because spatial parameters such as polar angle are explicitly modeled in pRF models, we were able to quantify and interpret these differences.

Our results have important implications for the study of memory reactivation. First, our findings suggest that the specific architecture of a sensory system may constrain what memory reactivation looks like in that system. Though memory reactivation is often studied in sensory domains, the architecture of these systems is not usually considered when interpreting reactivation effects. Here, we propose that hierarchical spatial pooling in visual cortex produces a systematic and distinct pattern of memory reactivation that cannot be attributed to retrieval failure or memory error. However, whether this architecture has any consequences for memory behavior is not clear from the present study. This question will be critical for future studies to address. Second, our results advocate for shifting away from the concept of memory reactivation as it has been understood and applied in the field of neuroimaging. Most previous work has focused on identifying *similarities* between the neural substrates of visual perception and visual memory. These studies have been successful in that they have produced many positive findings of memory reactivation in human visual cortex (Kosslyn et al., 1995; O’Craven & Kanwisher, 2000; Wheeler et al., 2000; Slotnick et al., 2005; Polyn et al., 2005; Kuhl et al., 2011; Bosch et al., 2014; Waldhauser et al., 2016; Lee et al., 2018; Bone et al., 2018). However, much of this work implicitly assumes that any mismatch between perception and memory is due to the fact that memory reactivation is either inherently low fidelity or susceptible to noise (Pearson et al., 2015), or is a subset of the perceptual response (Wheeler et al., 2000). Our results demonstrate that, at least in the spatial domain, this is not the case, and that systematic differences beyond noise exist. These results are broadly consistent with other recent findings suggesting computational differences between perception and memory derived from behavior (Bloem et al., 2018) and multivoxel pattern differences in perceived and remembered object representations measured with fMRI (Lee et al., 2012). Ultimately, the field should strive to identify, quantify, and explain these differences in order to fully understand the neural basis of memory. Using encoding models borrowed from sensory neuroscience to parameterize the differences between perception and memory may prove a fruitful way of making progress on this goal.

### Why do perceptual and mnemonic representations differ in visual cortex?

Despite the usefulness of encoding models like pRF models for quantifying neural responses in a stimulus-referred space, these models may not provide a natural explanation for *why* perception and memory responses differ. We show in Figure 6 that pRF models fail to capture the aspects of memory responses that are distinct from perceptual responses: namely, the dramatic change in spatial precision. While it would be possible to fit separate pRF parameters to memory data to improve the ability of the model to accurately predict memory responses, this still would not explain why these parameters or responses differ. How then can we account for this? We were particularly intrigued by the possibility that differences between memory and perception activity are a direct consequence of the direction of processing in hierarchically-organized cortex. Hierarchical structure and feedback processing are not typically directly simulated in a pRF model but there is considerable evidence to suggest these factors are of interest. Studies of anatomical connectivity provide evidence that the visual system is organized approximately hierarchically (Felleman & Essen, 1991; Barone et al., 2000; for other perspectives see Zeki, 2015; Hilgetag & Goulas, 2020), and that most connections within the visual system are reciprocal (Felleman & Essen, 1991). Studies also show that the hippocampus sits atop the highest stage of the visual hierarchy, with reciprocal connections to high-level visual regions via the medial temporal lobe cortex (Van Hoesen & Pandya, 1975; Felleman & Essen, 1991; Suzuki & Amaral, 1994). These observations make the prediction that initial drive from the hippocampus during memory retrieval should propagate backwards through the visual system. Neural recordings from the macaque (Naya et al., 2001) and human (Hindy et al., 2016; Linde-Domingo et al., 2019; Dijkstra et al., 2019), as well as computational modeling (Horikawa & Kamitani, 2017) support this idea.

Based on these observations and our hypothesis, we constructed a hierarchical network model in which we could simulate top-down activity. Though this model shares some features of hierarchical models of object recognition (Riesenhuber & Poggio, 1999; Serre et al., 2007), we emphasize that it is much simpler. Our model is entirely linear, its parameters are fixed *a priori* (not the result of training), and it encodes only one stimulus feature: space (Kay et al., 2013b). Critically, in contrast to pRF models, which express each region’s activity as a function of the stimulus, our model expresses each region’s activity as a function of the previous region’s activity (Fukushima, 1980; Riesenhuber & Poggio, 1999), and can therefore simulate both feedforward and feedback processes (Heeger, 2017). While highly simplified, the simulations we performed in this network captured the dominant features of our data, providing a parsimonious explanation for our observations. Interestingly, our simulations also indicate that some trends present in our data warrant further investigation. For instance, while we could not conclude that the earliest visual cortical areas had the least precise responses during memory (a *reversal* of the perception pattern), our simulations suggest that this effect should be present, albeit significantly weaker than in the feedforward direction. Future work should target this small effect with a sufficiently powered experiment.

Our simulations also raise interesting questions and predictions about the consequences of visual cortical architecture for cognition. First, why have a hierarchical architecture in which the detailed information present in early layers cannot be reactivated? The hierarchical organization of the visual system is thought to give rise to the low-level feature invariance required for object recognition (Riesenhuber & Poggio, 1999; Serre et al., 2007). Our results raise the possibility that the benefits of such an architecture for recognition outweigh the cost of reduced precision in top-down responses. Whether the extent of this tradeoff differs between healthy individuals or between healthy and neuropsychiatric populations, and what consequences this structure has for behavior, are interesting questions for future research. Second, how is it that humans have spatially precise memories if visual cortical responses do not reflect this? One possibility is that read-out mechanisms are not sensitive to all of the properties of mnemonic activity we measured. For instance, memory decisions could be driven exclusively by the neural population with the strongest response (e.g. those at the peak of the polar angle response functions). Another possibility is that regions without hierarchical structure do not exhibit these properties and reactivation in these other regions is preferentially used to guide memory-based behavior. These, and other possibilities should be directly explored in future work. Finally, our hierarchical simulations highlight the need to carefully separate the contribution of visual cortical architecture on reactivation from the effects of cognitive manipulations or effects occurring upstream of visual cortex (e.g. in the hippocampus).

### Relation to other forms of memory and attention

Sensory reactivation during long-term memory retrieval has parallels to sensory engagement in other forms of memory such as iconic memory and working memory. Nonetheless there may also be differences in the specific way that sensory circuits are used across these forms of memory. One critical factor may be how recently the sensory circuit was activated by a stimulus at the time of memory retrieval. In iconic memory studies, very detailed information can be retrieved if probed within a second of the sensory input (Sperling, 1960). In working memory studies, sensory activity is thought to be maintained by active mechanisms from stimulus encoding through a seconds-long delay. Using similar methods to the ones we use here (Sprague & Serences, 2013; Ester et al., 2013), many working memory studies have shown that early visual areas contain retinotopically specific signals throughout a delay period (Sprague & Serences, 2013; Sprague et al., 2014; Rahmati et al., 2017), paralleling our findings. In imagery studies, eye-specific circuits presumed to be in V1 can be re-engaged if there is a delay of 5 minutes or less from when the subject viewed stimuli through the same eye, but not if there is a delay of 10 minutes (Ishai & Sagi, 1995). Hippocampally-dependent memory retrieval is thought to be capable of engaging visual cortex at much longer delays. Given that the mechanism for engaging sensory cortex may differ across these different forms of memory, the question of how similar sensory activation is across these timescales remains an important open question. For example, shorter-term forms of memory might, in principle, cause more spatially-specific reactivation in early visual cortex than what we observed in long-term memory. Informal comparisons between our data and stimulus reconstructions made from working memory delay period activity (Sprague & Serences, 2013; Rahmati et al., 2017; Rademaker et al., 2019) suggest this may be the case, but a direct comparison is warranted. The current study offers a quantitative approach for directly comparing spatial tuning properties across different cognitive processes, and could be extended to include multiple forms of memory within the same experiment.

Are the spatial responses we observed during memory retrieval better characterized as long-term memory reactivation or as a special case of (memory-guided) spatial attention? Our results raise interesting questions about whether long-term spatial memory and endogenous spatial attention share mechanisms for modulating the response of visual cortical populations. In typical endogenous spatial attention tasks, subjects are explicitly cued to the most likely location of an upcoming stimulus prior to being presented with a difficult visual judgment (Carrasco, 2011). fMRI studies have repeatedly found that spatial attention enhances visually-evoked responses in visual cortex (Somers et al., 1999; Gandhi et al., 1999; Buracas & Boynton, 2007; Li et al., 2008). Similar to our results, spatial attention has also been shown to elicit spatially localized activation in the absence of any visual stimulation (Luck et al., 1997; Kastner et al., 1999; Chawla et al., 1999; Ress et al., 2000). It is at least logically possible for attention and memory to dissociate. Most endogenous attention tasks have no memory component since the cue explicitly represents the attended location. In contrast, in most episodic memory tasks the association between a cue and a stimulus is intentionally arbitrary so that it must be acquired and retrieved in a hippocampally-dependent manner. However, it is possible that spatial attention and memory processes only differ in their dependency on the hippocampus to retrieve the target location. Once this target location is determined, the same mechanisms could be used to initiate enhanced processing of the target location in sensory areas. Future experiments and modeling efforts should determine whether memory-driven and attention-driven activations in visual cortex differ, and whether it’s possible to develop a model of top-down processing in visual cortex that can account for both sets of observations.

## Conclusion

In the current work, we provide novel empirical evidence that memory retrieval elicits systematically different activation in human visual cortex compared to visual perception. Using simulations and a network model of cortex, we argue that these distinctions arise from a reversal of information flow within a hierarchically structured visual system. Collectively, this work makes progress on providing a detailed account of reactivation in visual cortex and sheds light on the broader computational principles that guide top-down processes in sensory systems.

## Methods

### Subjects

Nine human subjects participated in the experiment (5 males, 22–46 years old). All subjects had normal or correct-to-normal visual acuity, normal color vision, and no MRI contraindications. Subjects were recruited from the New York University community and included author S.E.F and author J.W. All subjects gave written informed consent to procedures approved by the New York University Institutional Review Board prior to participation. No subjects were excluded from data analysis.

### Stimuli

Experimental stimuli included nine unique radial frequency patterns (Fig. 1a). We first generated patterns that differed along two dimensions: radial frequency and amplitude. We chose stimuli that tiled a one dimensional subspace of this two dimensional space, with radial frequency inversely proportional to amplitude. The nine chosen stimuli took radial frequency and amplitude values of: [2, .9], [3, .8], [4, .7], [5, .6], [6, .5], [7, .4], [8, .3], [9, .2], [10, .1]. We selected four of these stimuli to train subjects on in the behavioral training session and to appear in the fMRI session. For every subject, those stimuli were: [3, .8], [5, .6], [7, .4], [9, .2]; (radial frequency, amplitude). The remaining five stimuli were used as lures in the test trials of the behavioral training session. Stimuli were saved as images and cropped to the same size.

### Experimental procedure

The experiment began with a behavioral training session, during which subjects learned four paired associates (Fig. 1). Specifically, subjects learned that four colored fixation dot cues were uniquely associated with four spatially localized radial frequency patterns. An fMRI session immediately followed completion of the behavioral session (Fig. 2a). During the scan, subjects participated in two types of functional runs (approximately 3.5 min each): (1) perception, where they viewed the cues and associated spatial stimuli; and (2) memory, where they viewed only the fixation cues and recalled the associated spatial stimuli. Details for each of these phases are described below. A separate retinotopic mapping session was also performed for each subject (Fig. 2b), which is described in the next section.

#### Behavioral training

For each subject, the four radial frequency patterns were first randomly assigned to one of four polar angle locations in the visual field (45°, 135°, 225°, or 315°) and to one of four colored cues (orange, magenta, blue, green; Fig. 1b). Immediately before scanning, subjects learned the association between the four colored cues and the four spatially localized stimuli through interleaved study and test blocks (Fig. 1c). Subjects alternated between study and test blocks, completing a minimum of four blocks of each type. Subjects were required to reach at least 95% accuracy, and performed additional rounds of study-test if they did not reach this threshold after four test blocks.

During study blocks, subjects were presented with the associations. Subjects were instructed to maintain central fixation and to learn each of the four associations in anticipation of a memory test. At the start of each study trial (Fig. 1c), a central white fixation dot (radius = 0.1 dva) switched to one of the four cue colors. After a 1 sec delay, the associated radial frequency pattern appeared at 2^°^ of eccentricity and its assigned polar angle location in the visual field. Each pattern image subtended 1.5 dva and was presented for 2 sec. The fixation dot then returned to white, and the next trial began after a 2 sec interval. No subject responses were required. Each study block contained 16 trials (4 trials per association), presented in random order.

During test blocks, subjects were presented with the colored fixation dot cues and tested on their memory for the associated stimulus pattern and spatial location. Subjects were instructed to maintain central fixation and to try to covertly recall each stimulus when cued, and then to respond to the test probe when prompted. At the start of each test trial (Fig. 1c), the central white fixation dot switched to one of the four cue colors. This cue remained on the screen for 2.5 sec while subjects attempted to covertly retrieve the associated stimulus. At the end of this period, a test stimulus was presented at 2° of eccentricity for 2 sec. Then, subjects were cued to make two consecutive responses to the test stimulus: whether it was the correct radial frequency pattern (yes/no) and whether it was presented at the correct polar angle location (yes/no). Each test stimulus had a 50% probability of being the correct pattern. Incorrect patterns were drawn randomly from the three patterns associated with other cues and the five lure patterns (Fig. 1a). Each test stimulus had a 50% probability of being in the correct polar angle location, which was independent from the probability of being the correct pattern. Incorrect polar angle locations were drawn from the three locations assigned to the other patterns and 20 other evenly spaced locations around the visual field (Fig. 1a). This placed the closest spatial lure at 15° of polar angle away from the correct location. Responses were solicited from the subject with the words “Correct pattern?” or “Correct location?” displayed centrally in white text. The order of these queries was counterbalanced across test blocks. Subjects responses were recorded on a keyboard with a maximum response window of 2 sec. Immediately after a response was made or the response window closed, the color of the text turned black to indicate an incorrect response if one was made. After this occurred for both queries, subjects were presented with the colored fixation dot cue and correct spatially localized pattern for 1 sec as feedback. This feedback occurred for every trial, regardless of subject responses to the probe. Each test block contained 16 trials (4 trials per association), presented in random order.

#### fMRI session

During the fMRI session, subjects participated in two types of functional runs: perception and memory retrieval (Fig. 2a). Subjects completed 5–6 runs each of perception and memory in an interleaved order. This amounted to 40–48 repetitions of perceiving each stimulus and of remembering each stimulus per subject.

During perception runs, subjects viewed the colored fixation dot cues and the radial frequency patterns in their learned locations. Subjects were instructed to maintain central fixation and to perform a one-back task on the stimuli. The purpose of the one-back task was to encourage covert stimulus-directed attention on each trial. At the start of each perception trial (Fig. 2a, top), a central white fixation dot (radius = 0.1 dva) switched to one of the four cue colors. After a 0.5 sec delay, the associated radial frequency pattern appeared at 2° of eccentricity and its assigned polar angle location in the visual field. Each pattern subtended 1.5 dva and was presented for 2.5 sec. The fixation dot then returned to white and the next trial began after a variable interval. Intervals were drawn from an approximately geometric distribution sampled at 3, 4, 5, and 6 sec with probabilities of 0.5625, 0.25, 0.125, and 0.0625 respectively. Subjects indicated when a stimulus repeated from the previous trial using a button box. Responses were accepted during the stimulus presentation or during the interstimulus interval. Each perception run contained 32 trials (8 trials per stimulus). The trial order was randomized for each run, separately for every subject.

During memory runs, subjects viewed the colored fixation dot cues and recalled the associated patterns in their learned spatial locations. Subjects were instructed to maintain central fixation, to use the cues to initiate recollection, and to make a subjective judgment about the vividness of their memory on each trial. The purpose of the vividness task was to enforce attention to the remembered stimulus on each trial. At the start of each memory trial (Fig. 2a, top), the central white fixation dot switched to one of the four cue colors. This cue remained on the screen for a recollection period of 3 sec. The fixation dot then returned to white and the next trial began after a variable interval. Subjects indicated whether the stimulus associated with the cue was vividly remembered, weakly remembered, or not remembered using a button box. Responses were accepted during the cue presentation or during the interstimulus interval. Each memory run contained 32 trials (8 trials per stimulus). For a given subject, each memory run’s trial order and trial onsets were exactly matched to one of the perception runs. The order of these matched memory runs was scrambled relative to the order of the perception runs.

### Retinotopic mapping procedure

Each subject completed either 6 or 12 identical retinotopic mapping runs in a separate fMRI session from the main experiment (Fig. 2b, top). Stimuli and procedures for the retinotopic mapping session were based on those used by the Human Connectome Project (Benson et al., 2018) and were identical to those reported in Benson & Winawer (2018). During each functional run, bar apertures on a uniform gray background swept across the central 24 degrees of the subject’s visual field (circular aperture with a radius of 12 dva). Bar apertures were a constant width (1.5 dva) at all eccentricities. Each sweep began at one of eight equally spaced positions around the edge of the circular aperture, oriented perpendicularly to the direction of the sweep. Horizontal and vertical sweeps traversed the entire diameter of the circular aperture while diagonal sweeps stopped halfway and were followed by a blank period. A full-field sweep or half-field sweep plus blank period took 24 s to complete. One functional run contained 8 sweeps, taking 192 s in total. Bar apertures contained a grayscale pink noise background with randomly placed faces, scenes, objects, and words at a variety of sizes. Noise background and stimuli were updated at a frequency of 3 Hz. Each run of the task had an identical design. Subjects were instructed to maintain fixation on a central dot and to use a button box to report whenever the dot changed color. Color changes occurred on average every 3 s.

### MRI acquisition

Images were acquired on a 3T Siemens Prisma MRI system at the Center for Brain Imaging at New York University. Functional images were acquired with a T2*-weighted multiband EPI sequence with whole-brain coverage (repetition time = 1 s, echo time = 37 ms, flip angle = 68°, 66 slices, 2×2×2 mm voxels, multiband acceleration factor = 6, phase-encoding = posterior-anterior) and a Siemens 64-channel head/neck coil. Spin echo images with anterior-posterior and posterior-anterior phase-encoding were collected to estimate the susceptibility-induced distortion present in the functional EPIs. Between one and three whole-brain T1-weighted MPRAGE 3D anatomical volumes (.8 x .8 x .8 mm voxels) were also acquired for seven subjects. For two subjects, previously acquired MPRAGE volumes (1×1×1 mm voxels) from a 3T Siemens Allegra head-only MRI system were used.

### MRI processing

#### Preprocessing

Anatomical and functional images were preprocessed using FSLv5.0.10 (Smith et al., 2004) and Freesurfer v5.3.0 (Fischl, 2012) tools implemented in a Nipype workflow (Gorgolewski et al., 2011). To correct for head motion, each functional image acquired in a session was realigned to a single band reference image and then registered to the spin echo distortion scan acquired with the same phase encoding direction. The two spin echo images with reversed phase encoding were used to estimate the susceptibility-induced distortion present in the EPIs. For each EPI volume, this nonlinear unwarping function was concatenated with the previous spatial registrations and applied with a single interpolation. Freesurfer was used to perform segmentation and cortical surface reconstruction on each subject’s average anatomical volume. Registration from the functional images to each subject’s anatomical volume was performed using boundary-based registration. Preprocessed functional time series were then projected onto each subject’s reconstructed cortical surface.

#### GLM analyses

Beginning with each subject’s surface-based time series, we used GLMdenoise (Kay et al., 2013a) to estimate the neural pattern of activity evoked by perceiving and remembering every stimulus (Fig. 2a). GLMdenoise improves signal-to-noise ratios in GLM analyses by identifying a pool of noise voxels whose responses are unrelated to the task and regressing them out of the time series. This technique first converts all time series to percent signal change and determines an optimal hemodynamic response function for all vertices using an iterative linear fitting procedure. It then identifies noise vertices as vertices with negative *R*^2^ values in the task-based model. Then, it derives noise regressors from the noise pool time series using principal components analysis and iteratively projects them out of the time series of all vertices, one noise regressor at a time. The optimal number of noise regressors is determined based on cross-validated *R*^2^ improvement for the task-based model. We estimated two models using this procedure. We constructed design matrices for the perception model to have four regressors of interest (one per stimulus), with events corresponding to stimulus presentation. Design matrices for the memory model were constructed the same way, with events corresponding to the the cued retrieval period. These models returned parameter estimates reflecting the BOLD amplitude evoked by perceiving or remembering a given stimulus versus baseline for every vertex on a subject’s cortical surface (Fig. 2a, bottom).

#### Fitting pRF models

Images from the retinotopic mapping session were preprocessed as above, but omitting the final step of projecting the time series to the cortical surface. Using these time series, nonlinear symmetric 2D Gaussian population receptive field (pRF) models were estimated in Vistasoft (Fig. 2b), as described previously (Dumoulin & Wandell, 2008; Kay et al., 2013b). We refer to this nonlinear version of the pRF model as the compressive spatial summation (CSS) model, following Kay et al. (2013b). Briefly, we estimated the receptive field parameters that, when applied to the drifting bar stimulus images, minimized the difference between the observed and predicted BOLD time series. First, stimulus images were converted to contrast apertures and downsampled to 101 x 101 grids. time series from each retinotopy run were resampled to anatomical space and restricted to gray matter voxels. time series were then averaged across runs. pRF models were solved using a two stage coarse-to-fine fit on the average time series. The first stage of the model fit was a coarse grid fit, which was used to find an approximate solution robust to local minima. This stage was solved on a volume-based time series that was first temporally decimated, spatially blurred on the cortical surface, and spatially subsampled. The parameters obtained with this fit were interpolated and then used as a seed for subsequent nonlinear optimization, or fine fit. This procedure yielded four final parameters of interest for every voxel: eccentricity (r), polar angle *(θ*), sigma (s), exponent (n). The eccentricity and polar angle parameters describe the location of the receptive field in space, the sigma parameter describes the size of the receptive field, and the exponent describes the amount of compressive spatial summation applied to responses from the receptive field. Eccentricity and polar angle parameters were converted from polar coordinates to rectangular coordinates (x, y) for some analyses. Variance explained by the pRF model with these parameters was also calculated for each voxel. All parameters were then projected from each subject’s anatomical volume to the cortical surface (Fig. 2b, bottom).

### ROI definitions

Regions of interest were defined by hand-drawing boundaries at polar angle reversals on each subject’s cortical surface, following established practice (Wandell et al., 2007). We used this method to define six ROIs spanning early to mid-level visual cortex: V1, V2, V3, hV4, LO (LO1 and LO2), and V3ab (V3a and V3b).

We further restricted each ROI by preferred eccentricity in order to isolate vertices responsive to our stimuli. We excluded vertices with eccentricity values less than 0.5° and greater than 8°. This procedure excluded vertices responding primarily to the fixation dot and vertices near the maximal extent of visual stimulation in the scanner. We also excluded vertices whose variance explained by the pRF model (*R*^2^) was less than 0.1, indicating poor spatial selectivity. All measures used to exclude vertices from ROIs were independent of the measurements made during the perception and memory tasks.

### Analyses quantifying perception and memory activity

Our main empirical analyses examined the evoked BOLD response to our experimental stimuli during perception and memory as a function of visual field parameters estimated from the pRF model. Our first step was to visualize evoked activity during perception and memory in visual field coordinates (Fig. 3a). Transforming the data in this way allowed us to view the activity in a common reference frame across all brain regions, rather than on the cortical surface, where comparisons are made difficult by the fact that surface area and cortical magnification differ substantially from one area to the next. To do this, we selected the (*x,y*) parameters for each surface vertex from the retinotopy model and the *β* parameters from the GLM analysis. Separately for a given ROI, subject, stimulus, and task (perception/memory), we interpolated the *β* values over (*x,y*) space. We rotated each of these representations according to the polar angle location of the stimulus so that they would be aligned at the upper vertical meridian. We then z-scored each representation before averaging across stimuli and subjects. We used these images to gain intuition about the response profiles and to guide subsequent quantitative analyses.

Before quantifying these representations, we simplified them further. Because our stimuli were all presented at the same eccentricity, we reduced our 2D stimulus coordinate representations to 1D dimensional responses functions on the polar angle dimension (Fig. 4a). We did this by selecting surface vertices whose (*x,y*) coordinates were within one s of the stimulus eccentricity (2°) for each ROI. We then binned the evoked BOLD response into 18 bins of polar angle distance from the stimulus and averaged within each bin to produce polar angle response functions for each subject. We divided each subject’s response function by the norm of the response vector before averaging across subjects and then multiplying by the average vector norm to get the correct units back. This procedure prevents a subject with a high BOLD response across all polar angles from dominating the average response. The resulting average polar angle response functions showed clear surround suppression for polar angles near the stimulus during perception. Given this, we fit a difference of two von Mises distributions to the average data, with the location parameters (*μ*) for the two von Mises distributions fixed to be equal, but the spread (κ) and scale allowed to differ.

We quantitatively assessed the similarities and differences between perception and memory responses using these fits. We examined the location parameter of the two von Mises distributions, and also computed the amplitude and FWHM of the fit. We repeated the fitting procedure 500 times, drawing subjects with replacement, to create bootstrapped 68% and 95% confidence intervals for both perception and memory location, amplitude, and FWHM parameters. To assess main effects of ROI, main effects of perception vs memory, and the interaction of these variables on location, amplitude, and FWHM values, we ran two-way ANOVAs. In all models, ROI was coded as an ordinal variable (V1 < V2 < V3 < hV4 < LO < V3ab) and perception/memory as a categorical variable. Because location, amplitude, and FWHM, were computed at the group-level and not at the single-subject level, we ran these ANOVAs using group-level values. We re-ran the ANOVAs for all 500 subject resamplings to create bootstrapped confidence intervals for ANOVA regression coefficients. We computed p-values for these effects by performing randomization tests. To create null distributions, we randomly shuffled the assignment of the location, amplitude, or FWHM values with respect to the independent variables of interest (ROI, perception/memory). We did this for every possible shuffling or a subset of 10,000 different shufflings, whichever was smaller. We then computed two-tailed p-values according to the position of the true regression coefficient in the null distribution. Statistical data visualizations for these analyses and those subsequently described were made using seaborn v0.9.0 (Waskom et al., 2018).

### Noise simulations

We performed three simulations designed to test whether differences in noise between perception and memory data could explain differences in the responses we observed. To this end, we identified three potential types of noise that were present in our memory data but not our perception data: 1) reduced SNR; 2) retrieval failure; 3) memory error. We then simulated the effect of these types of noise on our perception data and asked whether these noise sources could produce responses similar to the ones we observed during memory.

#### SNR simulation

To simulate reduced SNR, we created artificial datasets with different amounts of additive noise introduced to every vertex’s perception parameter estimate. Noise was added in five levels: noise needed to generate the empirical SNR of the perception data (p), noise needed to generate the empirical SNR of the memory data (m), or noise needed to generate 1/2, 1/4, or 1/8 the empirical SNR of the memory data. For each of these values, we simulated 100 independent datasets for every subject and ROI. We determined the amount of signal and noise actually observed for each vertex during perception and memory by examining bootstrapped parameter estimate distributions produced by GLMdenoise. We defined the median parameter estimate across bootstraps as the amount of signal and the standard error of this distribution as the amount of noise. To simulate new data for a vertex, we randomly drew a new parameter estimate from a normal distribution defined by the true signal value (median) and the noise value (SE) needed to produce the target SNR. Critically, we made the draws correlated across vertices for each simulation. We did this by selecting a scale factor from a standard normal distribution which determined how many SEs away from the median every vertex’s simulated value would lie. This scale factor was shared across all vertices in an ROI for a given simulation. This procedure overcompensates for the spatial correlation present in BOLD data by assuming that SNR is 100% correlated across all vertices in an ROI. Note that if the noise were uncorrelated across vertices, it would have a much smaller effect on the population tuning curves. For each noise value and each of the 100 simulations, we analyzed the simulated data using the same procedure we applied to the actual data. This yielded 100 von Mises fits to the simulated data for each noise value and ROI (Supplementary Fig. 1a). We extracted the location, amplitude, and FWHM values from these fits. We evaluated whether location and FWHM values approximated the ones we observed during memory by calculating the proportion of simulations that fell within the 95% confidence intervals derived from the memory data (Fig. 5a).

#### Retrieval failure simulation

To simulate retrieval failure, we created artificial datasets that contained a variable number of perception trials with no signal. Retrieval failure was simulated in five levels: 0%, 25%, 50%, 75%, and 100% of trials. For each of these values, we simulated 100 independent datasets for every subject and ROI. Depending on the level of retrieval failure, zero, one, two, three, or all four stimuli were randomly designated as ‘failed’ in each simulated dataset. For the failed stimuli, new parameter estimates were drawn from a distribution defined by zero signal during perception for every vertex. For the remaining stimuli, new parameter estimates were drawn from a distribution defined by the true perception signal for every vertex. Noise was equated for both trial types; for each vertex, we used the the amount of noise observed during perception. As in the SNR simulation, simulated data were correlated across vertices in an ROI and simulated data were analyzed using the same procedures as for the actual data. We evaluated whether simulated location and FWHM values approximated the ones we observed during memory by calculating the proportion of simulations that fell within the 95% confidence intervals derived from the memory data (Supplementary Fig. 1b and Fig. 5b).

#### Memory error simulation

To simulate memory error, we created artificial datasets that contained a variable amount of angular error in the peak location of the perception polar angle response functions. Memory error was simulated in seven levels of standard deviation: 0, 15, 30, 45, 60, 75, and 90 degrees. For each of these values, we simulated 100 independent datasets for every subject and ROI. We assigned the amount of memory error for a given subject and stimulus by drawing a random value from a normal distribution centered at the true angular location of the stimulus and with the current standard deviation. We then used these memory error values to misalign simulated perception data. Specifically, we created new perception datasets based on the *true* signal and noise characteristics of our perception data (equivalent to SNR simulation with ‘p’ noise or 0% retrieval failure simulation). As in prior simulations, simulated data were correlated across vertices in an ROI, and simulated data were analyzed according to the same procedure as for the actual data. Before averaging the simulated data across stimuli and subjects, we rotated each response by the chosen memory error value rather than by the location of that stimulus. That is, instead of rotating the response to a 45^°^ stimulus by 45^°^ to align all stimuli at 0^°^ (as we did in our main analysis), we rotated the response by a value either close (generating using small standard deviations, representing small errors) or potentially quite far away (generating using large standard deviations, representing large errors). After averaging, we extracted location and FWHM values. We then evaluated whether simulated location and FWHM values approximated the ones we observed during memory by calculating the proportion of simulations that fell within the 95% confidence intervals derived from the memory data (Supplementary Fig. 1c and Fig. 5c).

### pRF forward model

We evaluated the ability of our pRF model to account for our perception and memory measurements. To do this, we used our pRF model as a forward model. This means that we took the pRF model parameters fit to fMRI data from the retinotopy session (which used a drifting bar stimulus) and used them to generate predicted BOLD responses to our four experimental stimuli. The model takes processed stimulus images as input, and for each of these images, outputs a predicted BOLD response (in units of % signal change) for every cortical surface vertex. Before running the model, we transformed our experimental stimuli into binary contrast apertures with values of 1 where the stimulus was and values of 0 everywhere else. These images were downsampled to the same resolution as the images used to fit the pRF model (101 x 101).

#### Model specification

The pRF forward model has two fundamental operations. In the first operation, a stimulus contrast aperture image is multiplied by a voxel’s pRF. In the CSS and linear models, this pRF is defined as a circular symmetric 2D Gaussian, parameterized by a location in the visual field (*x,y*) and a size (*σ*). In the DoG+CSS version of the model, this pRF is defined as the difference of two such Gaussians, centered at the same location (see next paragraph). The second operation applies a power-law exponent (n) to the result of the multiplication, effectively boosting small responses. This nonlinear operation is the key component of the CSS model and improves model accuracy in high-level visual areas that are known to exhibit subadditive spatial summation (Kay et al., 2013b; Mackey et al., 2017). The values of the exponent range from 0 to 1, where a value of 1 returns the model to linear. The output of this nonlinear stage is multiplied by a final scale parameter (*β*), which returns the units to % signal change (Fig. 6a).

Because we observed negative surround responses in V1-V3 during perception, we focused mainly on the results of the DoG+CSS model. Prior work has shown that difference-of-Gaussians (DoG) pRF models can account for the center-surround structure we observed (Zuiderbaan et al., 2012). In order to construct DoG pRFs, we converted each pRF from the CSS model we fit to the retinotopy data to a DoG pRF. We chose this approach after encountering difficulty in fitting a DoG pRF model to the retinotopy data. First, we took every 2D Gaussian pRF from the CSS model, and we subtracted from it a second 2D Gaussian pRF that was centered at the same location but was twice as wide and half as high. This ratio of 2σ and .5*β* between the negative and positive Gaussians was fixed for all voxels. In order to prevent the resulting DoG pRF from being systematically narrower and lower in amplitude than the original pRF, we rescaled the σ and *β* of the original pRF before converting it to a DoG. We multiplied the original σ by 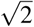 and the original *β* by 2, resulting in a DoG pRF with equivalent FWHM and amplitude as the original pRF. Thus, the DoG pRF differed from the original pRF only in the presence of a suppressive surround.

We compared the predicts of the DoG+CSS model to the results of the CSS model and to a linear model that we fit separately to the retinotopy data. In this linear model, no exponent parameter was fit. After generating a prediction for each subject, stimulus, and surface vertex, for each of our three forward models, we carried these predictions forward through the same analysis pipeline used to analyze our task-based data. This generated predicted polar angle response functions for each of the three pRF forward models (Fig. 6b and Supplementary Fig. 2). We generated bootstrapped predictions by conducting the same procedure on the bootstrapped datasets.

#### Evaluating model predictions

We next compared how well the DOG+CSS model predictions matched our perception versus memory measurements. We correlated the predicted location, amplitude, and FWHM parameters for each ROI with the actual perception and memory parameters. We evaluated these relationships by fitting a linear model to the predicted versus observed observations. To generate confidence intervals on these fits, we fit linear models to the 500 bootstrapped perception and memory datasets and the yoked pRF predictions (Fig. 6c).

We also compared the model accuracy of the DoG+CSS and CSS predictions alongside a linear prediction with no exponent parameter (Supplementary Fig. 2a). We calculated the coefficient of determination (*R*^2^) for the predicted polar angle response functions in each ROI, separately for the observed perception and memory polar angle response functions (Supplementary Fig. 2b). Under this measure, a model that predicts the mean observed response for every value of polar angle distance will have an *R*^2^ of zero, with better models producing positive values and worse models producing negative values. We generated confidence intervals for these accuracies by computing *R*^2^ values for each of the 500 bootstrapped perception and memory datasets and the yoked pRF predictions.

### Hierarchical network model

We assessed whether a simple instantiation of a single neural network model could account for both the perception and memory data. We implemented a fully linear hierarchical model of neocortex in which the activity from each layer was created by pooling activity from the previous layer. This model encodes 1D space only and its parameters are fixed (i.e. it is not trained). For the feedforward simulation, we began with a 1D square wave stimulus, which spanned −20 to 20 degrees of polar angle. We created a fixed Gaussian convolution kernel *(μ =* 0, s = 15), which we convolved with the stimulus to create the activity in layer 1. This layer 1 activity was convolved with the same Gaussian kernel to create the layer 2 activity, and this process was repeated recursively for 8 layers (Fig. 7a, left). In order to simulate memory-evoked responses in this network, we made two assumptions. First, we assumed that the feedback simulation began with the layer 8 activity from the feedforward simulation. That is, we assumed no information loss or distortion between perception and memory in the last layer. Second, we assumed that all connections were reciprocal and thus that the same Gaussian kernel was applied to transform layers in the feedback direction as in the feedforward direction (Fig. 7a, right). Thus, in the feedback simulation, we convolved the layer 8 activity with the Gaussian kernel to produce the layer 7 activity and repeated this procedure recursively, ending at layer 1 (Fig. 7b). Note that these computations can be performed with matrix multiplication rather than convolution by converting the convolutional kernel to a Toeplitz matrix, which is how we implemented it. In this case, the transpose of the Toeplitz matrix (itself, as it is symmetric) is used in the feedback direction. We plot the location, amplitude and FWHM for each layer’s activation in the same convention as the data (Fig. 7c).

## Acknowledgements

S.E.F. was supported by an NIH Blueprint D-SPAN F99/K00 Award (F99-NS105223).

## Author Contributions

Conceptualization, S.E.F., B.A.K., and J.W.; Methodology, S.E.F. and J.W.; Software, S.E.F. and J.W.; Investigation, S.E.F.; Writing–Original Draft, S.E.F. and J.W.; Writing–Review & Editing, S.E.F., B.A.K., and J.W.; Funding Acquisition, S.E.F.; Supervision, B.A.K and J.W.

## Supplementary Figures

**Figure 1.**
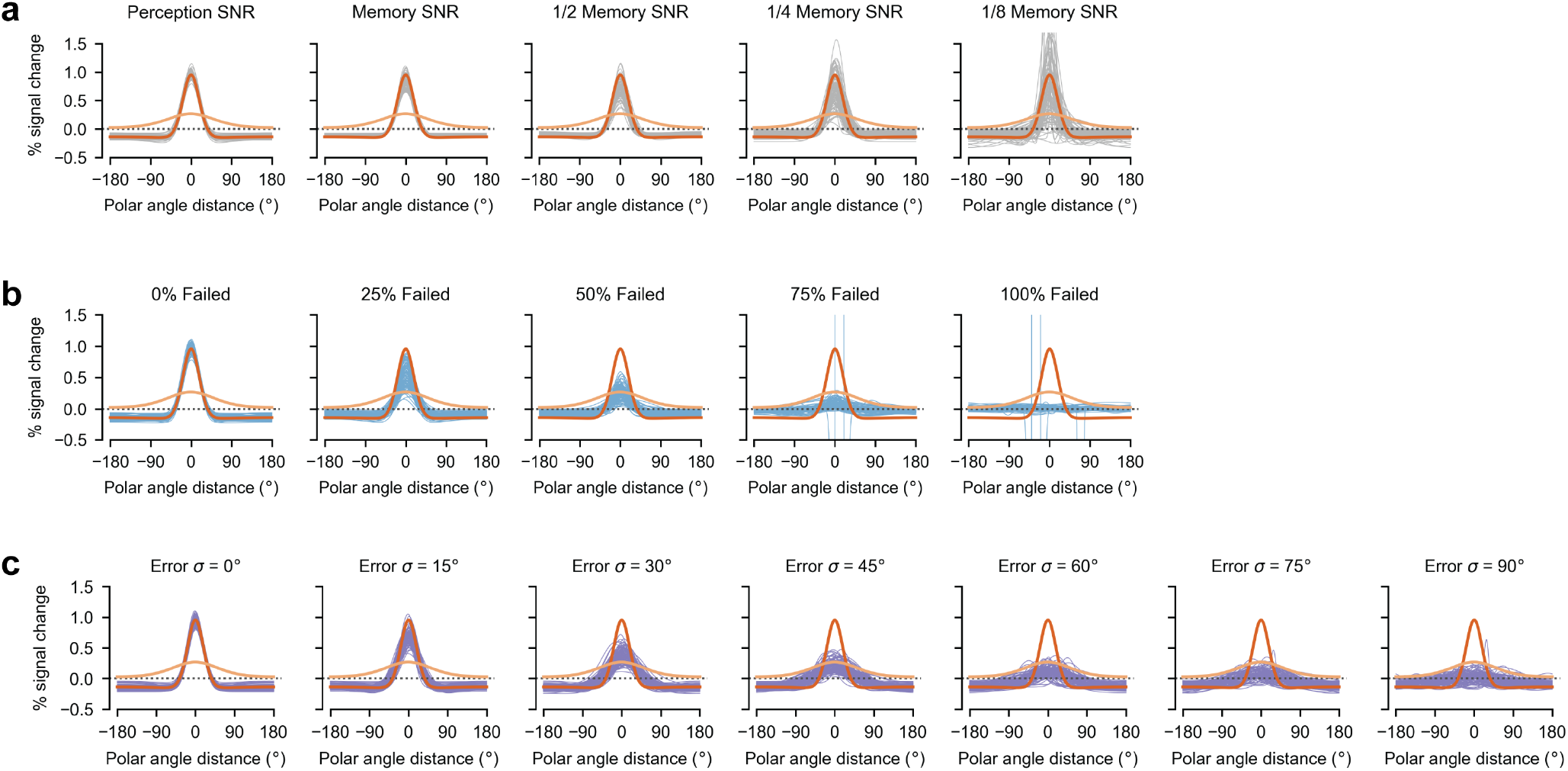
Simulated V1 datasets with different noise levels. **(a)** Gray lines represent the fits to simulated V1 perception datasets with different levels of SNR. Each panel contains 100 independently simulated datasets with the same noise level. Orange lines represent the fits to the actual perception and memory data, reproduced from Figure 4b, and are the same for each SNR value. **(b)** Purple lines represent the fits to simulated V1 perception datasets with different frequencies of failed retrieval. Other conventions as in (a). **(c)** Blues lines represent the fits to simulated V1 perception datasets with different amounts of memory error. Other conventions as in (a).

**Figure 2.**
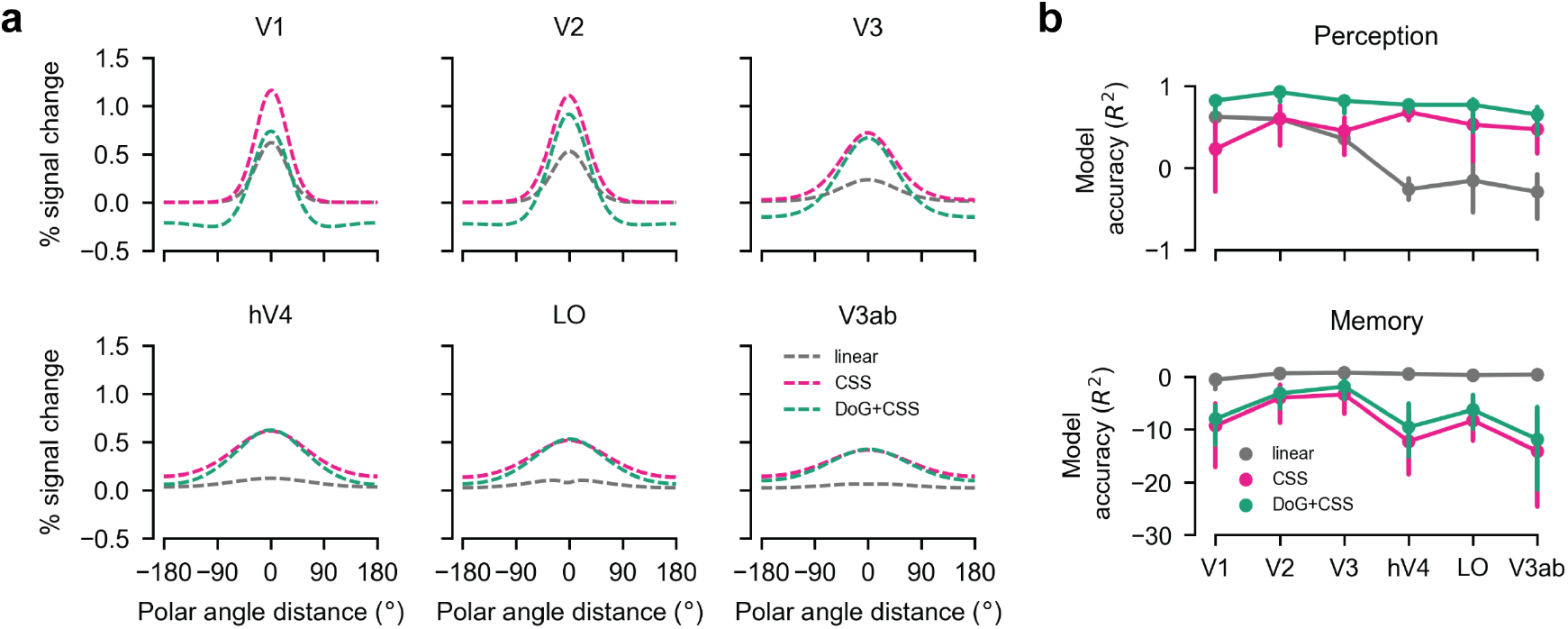
pRF model comparisons. **(a)** Predicted polar angle response functions are plotted for three pRF models: linear, CSS, and DoG+CSS. Comparing these responses to perception data plotted in Figure 4b, the linear model did the poorest job of predicting perception responses. Linear predictions underestimated the amplitude of the observed response, particularly in later visual areas. Both nonlinear models (CSS and DOG+CSS) avoided this magnitude of failure. The DoG+CSS model selectively captured negative responses in V1–V3. **(b)** Model accuracy (*R*^2^) of the predicted polar angle response functions for each pRF model, evaluated separately for perception and memory data in each ROI. Error bars indicate 68% bootstrapped confidence intervals. Accuracy of the linear model in predicting perception data dropped steadily moving away from V1, indicating poor fit. Model accuracies for the the CSS and DoG+CSS models were higher and more stable across ROIs, with the DoG+CSS performing slightly better in every region. With the exception of the linear model in late visual areas, accuracy for all three models was far worse for memory data than perception data.

